# A ribozyme ligase that requires a 3′ terminal phosphate on its RNA substrate

**DOI:** 10.1101/2025.05.06.652512

**Authors:** Annyesha Biswas, Zoe Weiss, Jack W. Szostak, Saurja DasGupta

**Author notes:** Corresponding Author Jack W. Szostak − Howard Hughes Medical Institute, Department of Chemistry, University of Chicago, Chicago, Illinois, 60637, USA. Corresponding Author Saurja DasGupta − Department of Chemistry and Biochemistry, University of Notre Dame, Notre Dame, Indiana, 46556, USA. Department of Biological Sciences, University of Notre Dame, Notre Dame, Indiana, 46556, USA.

## Abstract

Ribozymes played essential roles in catalyzing metabolic processes and facilitating genome replication in primordial RNA-based life. In vitro evolution has enabled the discovery of ribozymes that catalyze diverse chemical reactions, but it is likely that many catalytic activities of RNA remain to be uncovered. Expanding our understanding of the biochemical capabilities of RNA, especially new ribozyme functions related to RNA assembly, will help to refine models of primordial genome replication and, more broadly, the origin and evolution of early life. Here, we report the discovery of a novel ribozyme ligase that catalyzes the attack of the 2′-hydroxyl group of an RNA substrate on the 5′-triphosphate of a second RNA, but only when the substrate RNA possesses a 3′-phosphate vicinal to its nucleophilic 2′-hydroxyl group. This activity emerged unexpectedly during a directed evolution experiment designed to isolate ribozyme ligases that use 5′-triphosphorylated RNA oligonucleotides as substrates. Because the 3′-phosphate group on the RNA substrate does not directly participate in the ligation reaction, it would be extremely challenging to design a selection strategy to isolate ribozyme ligases with this unique reactivity. The ligase’s requirement for an RNA 3′-phosphate group on the substrate resembles enzymatic mechanisms found in protein-based RNA repair pathways. We propose that ribozyme-catalyzed ligation of 3′-phosphorylated RNA could have enabled the repair of cleaved RNA strands in primordial cells. We demonstrate that our in vitro evolved ribozymes ligate themselves specifically to 3′-phosphorylated RNA fragments present in heterogenous mixtures of cellular RNA, demonstrating their potential utility as enrichment reagents for profiling RNA cleavage products in transcriptomic studies. Our findings not only report a new catalytic reactivity in RNA but also provide insights into ribozyme evolution, primordial RNA repair, and potential applications in RNA sequencing.

## INTRODUCTION

Enzyme catalysis is a fundamental feature of extant life. The emergence of catalytic properties in biomolecular scaffolds is central to one of the biggest mysteries in science – the origin of life – because the appearance and evolutionary diversification of the earliest enzymes likely determined the fate of primordial cell lineages. Early life is thought to have used enzymes made of RNA, referred to as ribozymes, to catalyze its metabolic reactions as well as to replicate its genomes, also made of RNA. Therefore, exploring the catalytic capabilities of ribozymes provides an opportunity to understand the origins of biocatalysis. The study of RNA catalysis has been greatly expanded by our ability to evolve ribozymes outside the cell.^1, 2^ In vitro selection from completely random RNA sequence libraries has yielded ribozyme reactivities that do not exist in nature and directed evolution of existing ribozymes has resulted in the isolation of new ribozymes with different and sometimes unexpected functions.^3^ Although in vitro selection protocols employ controlled selection pressures to isolate ribozymes with desired functions, it is not uncommon for ribozymes with unexpected functions to emerge from such experiments.^4, 5^ In those cases, ribozymes with unexpected activities use alternative (and often unforeseen) reaction pathways to become enriched through a selection protocol designed to yield a different desired activity. More than thirty years of artificial ribozyme evolution experiments have revealed that RNA can support a diversity of catalytic phenotypes.^3^ However, considering the small number of ribozyme selection experiments that have been carried out to date compared to the range of interesting metabolic and replication-related reactions that could potentially be catalyzed by RNA, it is clear that we have not yet approached the full catalytic potential of RNA.

The diversity of ribozyme activities is well illustrated by RNA ligases. These ribozymes catalyze ligation reactions between the activated 5′-phosphate of an RNA substrate and either the 2′-hydroxyl, 3′-hydroxyl, or 5′-phosphate of a second RNA substrate.^6–8^ Some ligases require an external template, while others do not.^6, 7^ Although certain ligase ribozymes function as true enzymes by catalyzing ligation between two separate RNA oligonucleotides in trans, the majority facilitate the ligation of an oligonucleotide substrate to itself.^6, 7^ The many variations of a simple RNA ligase phenotype further highlight the possibility of discovering new but related ribozyme reactivities. In this work, we report the serendipitous discovery of a new ligase activity, where the ribozyme catalyzes a reaction between its 5′-triphosphate group and the 2′-hydroxyl group of an RNA oligonucleotide substrate only when the substrate possesses a 3′-phosphate group vicinal to its 2′-hydroxyl nucleophile. Ribozymes with this activity were isolated from an in vitro evolution experiment designed to evolve ribozymes that catalyze ligation with 5′-triphosphorylated RNA substrates. It is interesting that of the isolated sequences, >60% exhibited this unexpected ligase activity, while only ∼30% exhibited the desired ‘triphosphate ligase’ activity via the generation of a canonical 3′-5′ phosphodiester linkage between the ribozyme and the substrate.

The absolute requirement for a 3′-phosphate on the RNA substrate, together with ligation through the 2′-hydroxyl, represents a surprising variation of the standard ribozyme ligase reactivity. Curiously, this reactivity is reminiscent of protein-based RNA ligase enzymes in extant biology that perform RNA repair in tRNA maturation pathways and in response to cellular stress.^9^ In that context, we outline plausible roles for such ribozymes in genome repair in the extinct biology of the RNA World. The unique specificity of the isolated ribozymes toward 3′-phosphorylated RNAs opens the possibility of adapting these ribozymes as reagents for the specific enrichment of cleaved RNAs from cells. To that end, we show that RNA cleavage products containing 3′-phosphorylated ends can be selectively captured from cellular RNA and amplified using these ribozymes. In addition to reporting a novel enzyme reactivity, our work demonstrates that a single selection pressure can lead to the enrichment of multiple catalytic functions and highlights the versatility of RNA as a catalytic molecule.

## RESULTS

### New ligase ribozymes isolated from in vitro evolution

Recently, we used in vitro evolution to ask whether and how ligase ribozymes might switch substrate specificity from RNA oligonucleotides activated with a 5′-phosphorimidazole group (5′-PI) to those containing the biologically relevant 5′-triphosphate group (5′-PPP).^10^ In that work, we created a partially randomized RNA library derived from a previously characterized ‘phosphorimidazolide ligase’ (AI-Ligase)^11^ by mutagenizing a 40 nt region of this ribozyme at 21% per nucleotide position. This variable region was flanked by constant regions, which provide binding sites for reverse transcription and PCR primers. The 3′ constant region included an 8 nt ‘primer’ sequence connected to the rest of the construct by a hexauridine linker. The terminal nucleotide of this primer sequence (i.e., the last nucleotide of the selection construct) was the intended site of ligation (Fig. 1A). Library sequences with the ability to ligate to a 16 nt RNA substrate containing a 5′-triphosphate group and 3′-biotin were purified away from unreactive sequences by streptavidin bead capture and subsequently amplified by RT-PCR and in vitro transcription (Fig. 1). The substrate sequence was connected to a triethylene glycol (TEG) linker by its 3′-phosphate group, and the TEG linker was in turn covalently attached to a biotin group. The target reaction for this selection involved a nucleophilic attack of the 3′-hydroxyl (3′-OH) end of the selection construct (the ‘primer’) on the α-phosphate of the 5′-PPP group of the substrate. (Fig. 1). Although we isolated five distinct classes of ligase ribozymes from that experiment, only one ribozyme class was found to catalyze the desired reaction.^10^ Here, we report that the other ribozyme classes that collectively covered >60% of the isolated RNA population catalyze a new and unexpected variation on this standard mode of RNA ligation.

**Figure 1.**
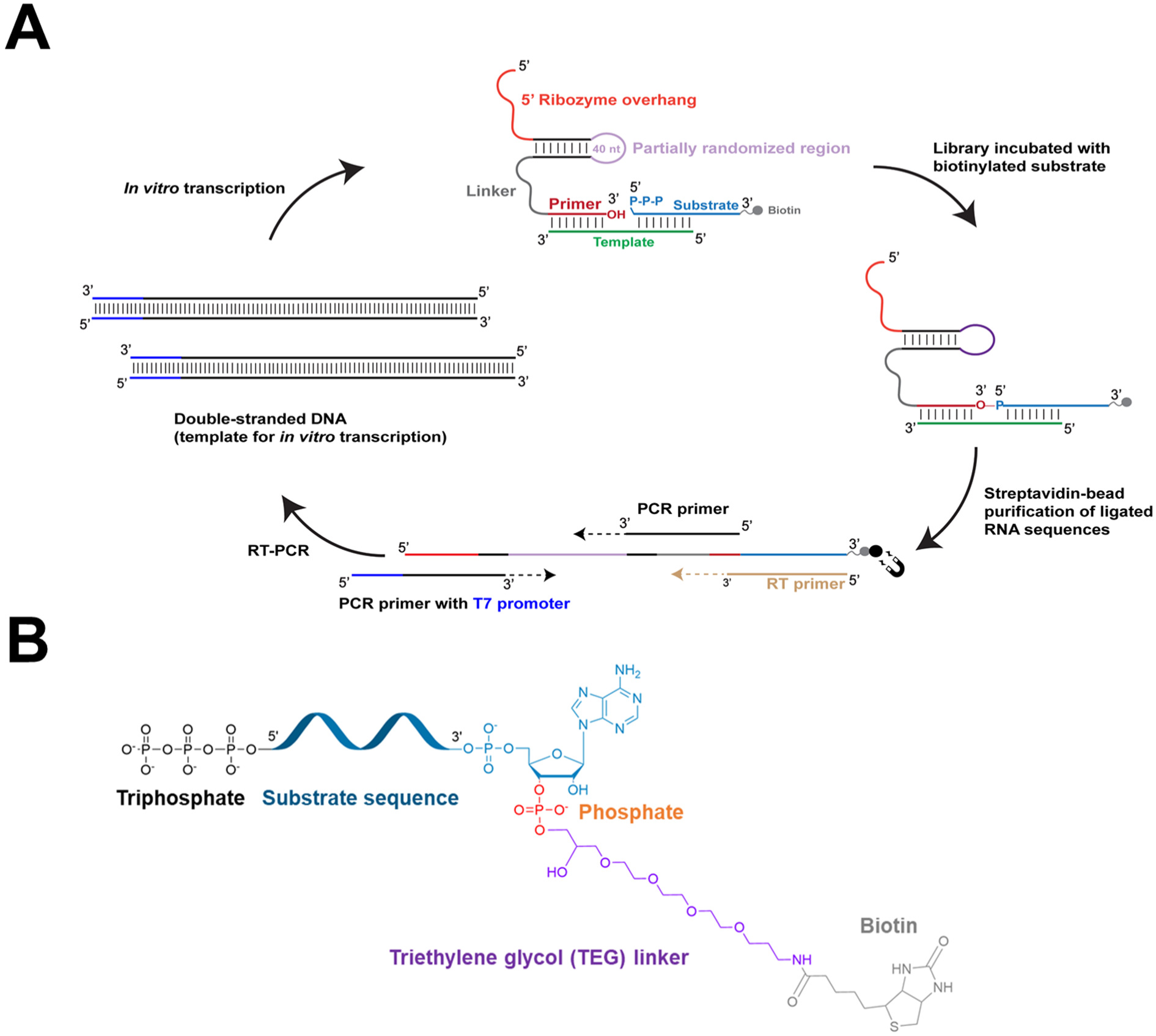
Selection protocol and substrate used to isolate ligase ribozymes. **A.** An RNA library containing a partially randomized sequence (depicted in light pink) derived from an AI-Ligase^11^ was challenged with an RNA substrate (depicted in light blue), containing a 5′-triphosphate group and a 3′-TEG-biotin group (depicted in gray) in the presence of an RNA template (depicted in green). Ligated sequences were purified by binding to streptavidin-coated magnetic beads and reverse transcribed using a primer (RT primer; depicted in gold) that is complementary to the entire substrate sequence. The RT primer also has the potential to bind directly to the library sequences by forming four base-pairs with the 3′ end of the ‘primer’, depicted in red. The cDNA was PCR-amplified, with the T7 promoter sequence (depicted in blue) added to the dsDNA sequence during PCR. This dsDNA was transcribed to generate the library for subsequent rounds of selection. **B.** The chemical features of the substrate used in the selection (PPP-Substrate-Biot). The substrate contains a triphosphate group (black) on its 5′ end and is connected to a biotin moiety (gray) via a triethylene glycol (TEG) linker (purple) attached to the 3′-phosphate (orange) on the terminal adenine of the substrate.

Outputs from each round were analyzed by high-throughput sequencing.^12^ Closely-related sequences isolated from the last round (round 6) were binned into clusters. We tested the peak sequences of the five most abundant clusters, referred to henceforth as CS1-CS5 (Extended Data Fig. 1), for their capacity to ligate to the 5′-triphosphorylated 3′-biotinylated, substrate (henceforth, PPP-Substrate-Biot) used in each selection round (Fig. 1B). Collectively, these sequences represented >90% of the entire selected population. CS1, CS2, CS4, and CS5 catalyzed ligation with rates between 0.6 h^-1^ to 1.5 h^-1^, yielding between 30% to 55% ligated product after 3 h. In contrast, CS3 showed reduced activity with a ∼75-fold lower ligation rate than CS1 (Fig. 2A, C, D). Interestingly, only CS3 catalyzed the desired ligation reaction with a PPP-Substrate, which we previously reported as a *bona fide* triphosphate ligase ribozyme.^10^ CS1, CS2, CS4, and CS5, on the other hand, appeared to exhibit an unexpected dependence on the 3′ biotinylation state of the substrate (Fig. 2B). This indicated that CS1, CS2, CS4, and CS5 do not catalyze the desired reaction. However, the appearance of a ligated product when incubated with PPP-Substrate-Biot suggests that these sequences are ligases that use an unforeseen reaction pathway. In the following sections, we investigate the unexpected features of these ligases and uncover a novel enzyme reactivity.

**Figure 2.**
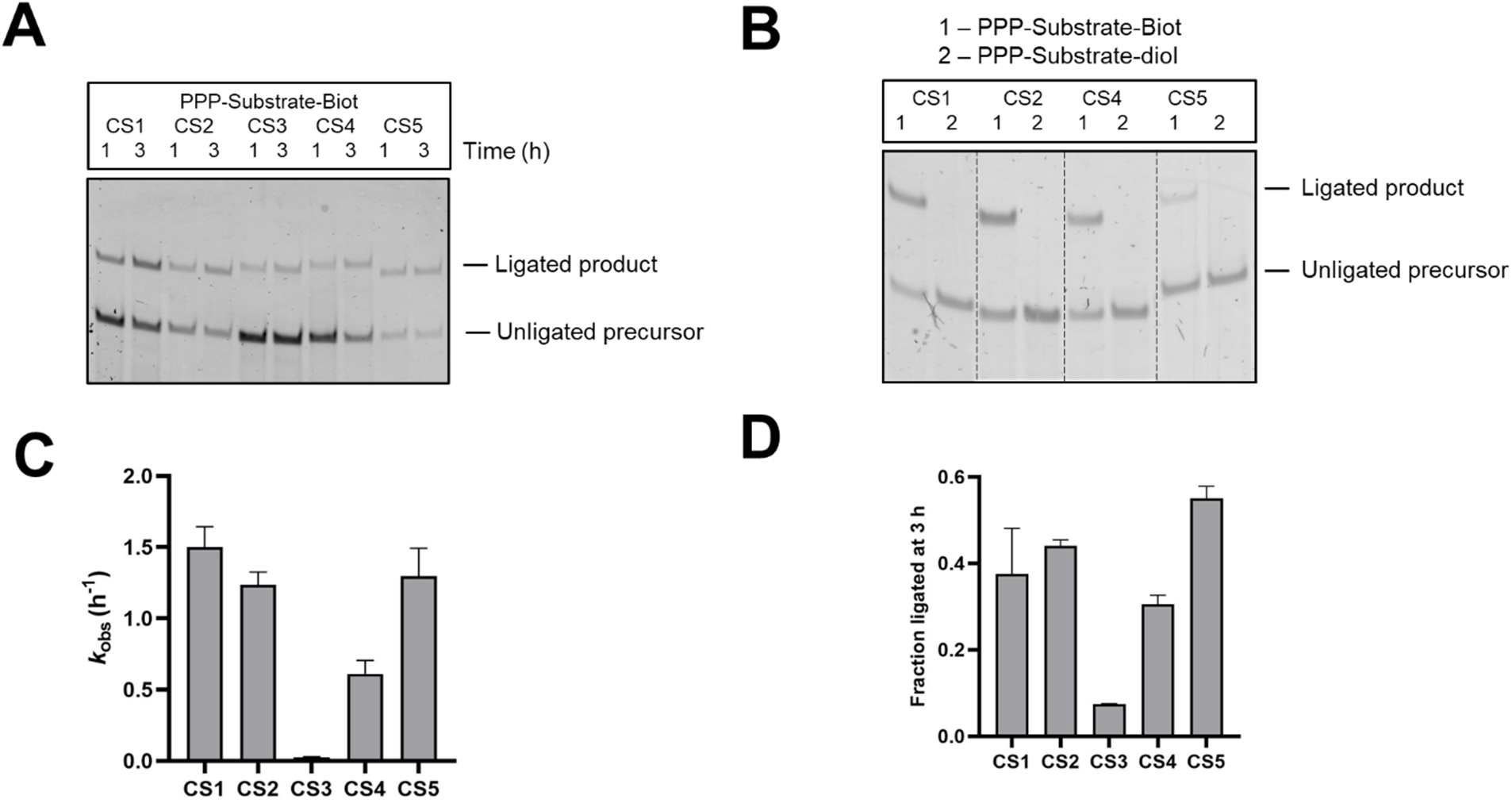
Ligase activities of the isolated ribozymes. **A.** Peak sequences from clusters 1-5, CS1-CS5, catalyze ligation with a substrate containing a 5′-triphosphate group and 3′-TEG-biotin group (PPP-Substrate-Biot). **B.** CS1, CS2, CS4, and CS5 do not ligate to a substrate oligonucleotide with a 2′,3′ cis-diol. **C.** CS1, CS2, CS4, and CS5 exhibit *k*_obs_ values of 0.6-1.5 h^-1^, but CS3-catalyzed ligation is significantly slower with PPP-Substrate-Biot. **D.** CS1, CS2, CS4, and CS5 ligated to 30-55%, while CS3 ligated to ∼7% with PPP-Substrate-Biot in 3 h. Ligation reactions contained 1 µM ribozyme, 1.2 µM RNA template, and 2 µM RNA substrate, PPP-Substrate-Biot, in 100 mM Tris-HCl (pH 8.0), 300 mM NaCl, and 100 mM MgCl_2_.

### Unexpected ligation to the ribozyme 5′ end

Given that CS1, CS2, CS4, and CS5 all exhibited an identical apparent dependence on the 3′ biotinylation state of the substrate, we selected the most abundant sequence, CS1, for detailed biochemical characterization. The desired reaction was a templated ligation between the 3′ and 5′ termini of the ribozyme and substrate, respectively, where the 16-nt template oligonucleotide was expected to bring the two RNA termini together by forming 8 base-pairs with sequences at the 3′ and 5′ end of each RNA (Fig. 1A). However, CS1-catalyzed ligation was only 3-fold slower in the absence of the external template (Fig. 3A, B). This absence of template requirement was reminiscent of a previous experiment aimed at the evolution of a ribozyme that catalyzes ligation between its 3′-OH and the 5′ end of an imidazole-activated RNA substrate.^8^ In that work, Chapman and Szostak unintentionally selected ribozymes that catalyze a ligation reaction between the ribozyme 5′-PPP (present because the ribozyme was generated by in vitro transcription) and the 5′-phosphorimidazole moiety of the substrate, generating a 5′-5′ linkage. To test the possibility that we had accidentally selected similar 5′-5′ ligases, we synthesized truncated versions of CS1 by deleting either its first 25 nucleotides (5′ truncation or 5′t) or its last 14 nucleotides (3′ truncation or 3′t) and tested for ligase activity. While the 3′ truncation preserved ligation, the 5′ truncated ribozyme was inactive for ligation (Fig. 3C). This suggested that the deleted 25 nt sequence at the ribozyme 5′ end is important for its activity, either by being part of its active fold or by directly participating in ligation. To decouple its potential structural vs. functional roles, we tested the activities of CS1-variants containing either a triphosphate (PPP), monophosphate (P), or hydroxyl (OH) group at their 5′ ends against substrate variants with either a 5′-PPP, 5′-P, or 5′-OH. Surprisingly, all substrate variants could be ligated to CS1, but only when CS1 itself contained a 5′-PPP group (Fig. 3D). Efficient ligation with 5′-P or 5′-OH substrates discounted the possibility of a nucleophilic attack by the ribozyme 5′-PPP on the substrate 5′-end (as observed by Chapman and Szostak)^8^. The inactivity of CS1 without a 5′-PPP, on the other hand, indicated that the ribozyme 5′ end was the likely site of attack by a nucleophilic group on the substrate.

**Figure 3.**
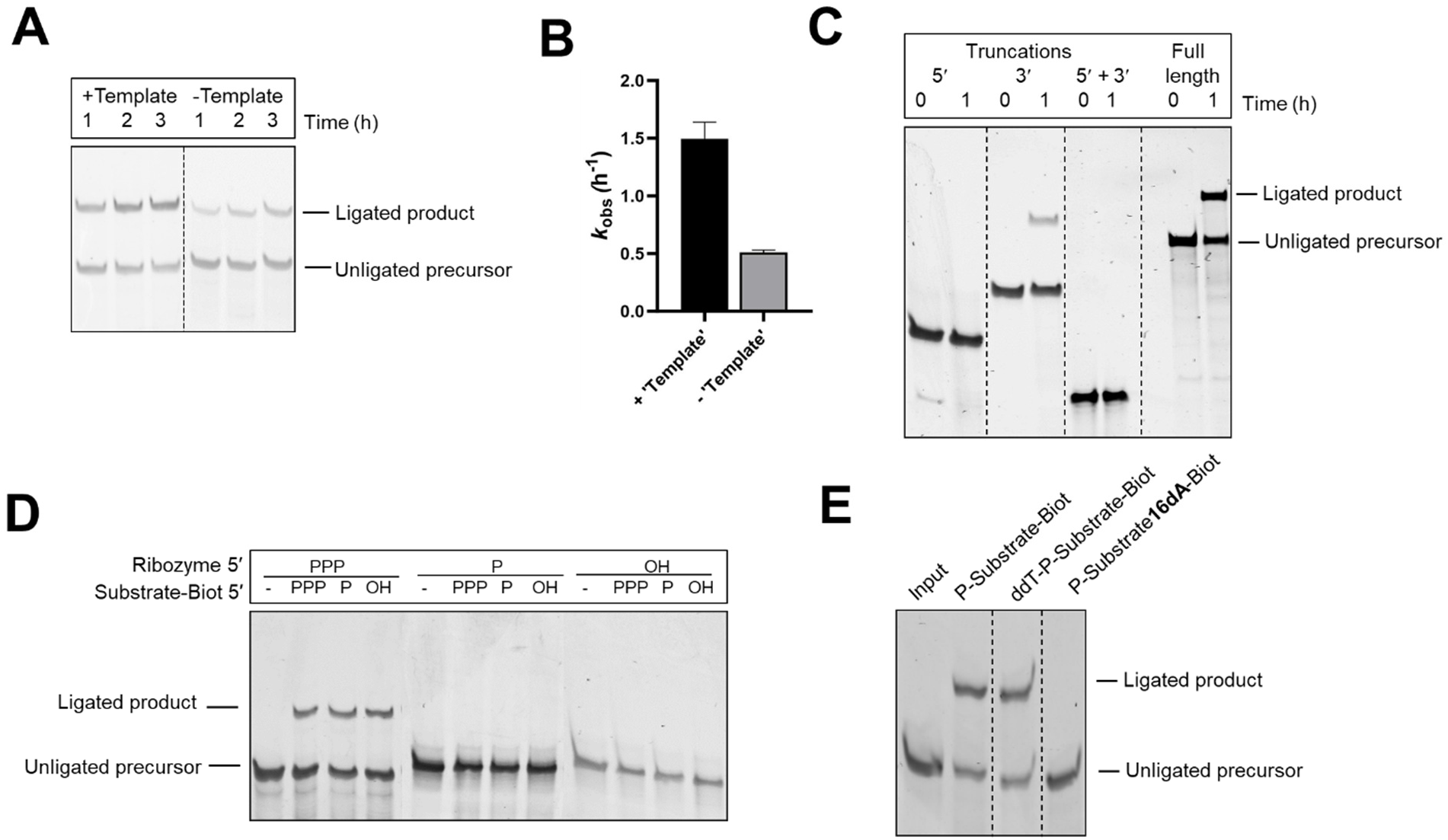
Untemplated ligation between the 5′-triphosphate group of CS1 and the 2′-hydroxyl group of the substrate. **A, B.** CS1-catalyzed ligation is preserved in the absence of an external template with only a 3-fold decrease in activity. **C.** Truncating CS1 by deleting its first 25 nucleotides abrogates ligation; however, deleting its last 14 nucleotides preserves activity. **D.** Ligation requires a triphosphate group at the ribozyme 5′ end but is agnostic to the chemistry at the substrate 5′ end. **E.** Blocking the 5′ end of the substrate with an inverted dideoxythymidine group (ddT-P-Substrate-Biot) preserves ligation; however, a substrate with a 3′ terminal deoxyribonucleotide (P-Substrate-**16dA** Biot) fails to ligate to CS1. Ligation reactions contained 1 µM ribozyme and 2 µM RNA substrate (PPP-Substrate-Biot unless otherwise specified) in 100 mM Tris-HCl (pH 8.0), 300 mM NaCl, and 100 mM MgCl_2_. Reactions do not contain an external template unless indicated (A) at 1.2 µM concentration.

### Reaction between the substrate 2′-hydroxyl and the ribozyme 5′-triphosphate groups

With the ribozyme 5′-PPP identified as the site of nucleophilic attack, we sought to uncover the corresponding nucleophile on the substrate. In principle, a highly promiscuous ribozyme ligase could catalyze the nucleophilic attack of substrate 5′-PPP, 5′-P, or 5′-OH groups on its own 5′-α-phosphate, with the release of a pyrophosphate group.^5, 13, 14^ However similar reactivities of all three substrates, under a range of different conditions, strongly suggested that the 5′ end of the substrate does not participate in ligation. CS1 exhibited similar reaction rates (*k*_obs_ = ∼1.5/h) with all three 5′-modified substrates (Extended Data Fig. 2) and exhibited comparable Mg^2+^ requirements with substrates possessing 5′-PPP or 5′-OH groups: [Mg^2+^]_1/2_ (HO-Substrate-Biot) = ∼15 mM; [Mg^2+^]_1/2_ (PPP-Substrate-Biot) = ∼20 mM (Extended Data Fig. 3). Additionally, CS1 showed similar pH-rate profiles (log *k*_obs_ vs pH) when ligating 5′-P or 5′-OH substrates, exhibiting linearity with a slope of ∼1 between pH 6.5 and 8 for both reactions (Extended Data Fig. 4). This usually indicates a single H^+^ transfer in the rate-determining step and is a feature of nucleophilic hydroxyl groups.^15^ The irrelevance of the substrate 5′ chemistry was conclusively shown by the ligation of CS1 to a substrate variant containing an inverted dideoxythymidine(ddT) group blocking its 5′ end (Fig. 3E).

With the 5′ end of the substrate discounted as a reactive center and the 3′ end of the substrate being blocked by TEG-biotin, we turned to the 2′-OH of the terminal adenine residue of the substrate as the most likely candidate for the nucleophile. A substrate where the terminal adenine ribonucleotide (rA16) was replaced by a deoxyribonucleotide (dA16) was inactive for ligation, confirming the substrate 2′-OH as the most likely nucleophile (Fig. 3E). Identical results with CS2, CS4, and CS5 established a common reactivity for all four ligases (Extended Data Fig. 5). Nucleophilic attack of the substrate 2′-OH on the ribozyme 5′-PPP would generate a noncanonical 2′-5′ linkage between the ribozyme and substrate. We performed a dual exonuclease digestion assay to test for the presence of this noncanonical 2′-5′ linkage. We purified the products of ligation between CS1 and substrates with either 5′-PPP or 5′-P groups and subjected them to 5′-P-dependent Terminator 5′◊3′ exonuclease digestion. As expected, the 5′-PPP ligated product was resistant to degradation (negative control), but the digestion of the 5′-P ligated product yielded RNA that was comparable in size to the ribozyme (Extended Data Fig. 6A). On the other hand, 3′◊5′ RNase R digestion of the ligated product obtained from the reaction with CS1 and a 5′-OH substrate generated RNA that was one nucleotide longer than the substrate (Extended Data Fig. 6B). These results suggest the presence of a 2′-5′ phosphodiester bond between the ribozyme and substrate and are consistent with the reaction between substrate terminal 2′-OH and ribozyme 5′-PPP groups.

The reactivity described above also explains the higher-order ligation products generated during CS1-catalyzed ligation with a 5′ 2AI-activated substrate (AIP-Substrate-Biot) in the presence of the external template and 100 mM Mg^2+^ (Extended Data Fig. 7). These higher-order products are concatemeric RNA sequences with a heterogenous backbone composed of both 2′-5′ and 3′-5′ phosphodiester linkages formed as a result of a combination of ribozyme-catalyzed 2′-5′ ligation and template-directed nonenzymatic 3′-5′ ligation at a high Mg^2+^ concentration. The production of concatemeric RNA demonstrates the ribozyme’s ability to function in the context of longer transcripts.

### Ligation requires a 3′-phosphate group on the substrate

The unexpected observation that substrates without the 3′-‘TEG-biotin’ moiety (i.e., with a 2′,3′ diol) are not substrates for ligation catalyzed by CS1, CS2, CS4, and CS5 (Fig. 2B) pointed to a role for the 3′-biotin in this reaction. As ribozymes that utilize thiamin (Vitamin B1) as a cofactor have been reported,^16^ we wondered if CS1, CS2, CS4, and CS5 might utilize biotin (Vitamin B7) for catalysis. The first indications that these ribozymes did not use biotin in their reactions came from the observation that supplementing a reaction between CS1 and an unbiotinylated substrate with free biotin does not rescue ligation (Fig. 4A). Moving the biotin modification from the 3′ to the 5′ end of the substrate eliminated ligation, underscoring the importance of a 3′ modification on the substrate. Further, CS1-catalyzed ligation was preserved on replacing the biotin moiety with desthiobiotin or deleting the TEG spacer between the substrate 3′-phosphate and the biotin (so that the 3′-phosphate was directly connected to the biotin) (Fig. 1B, Fig. 4B). Finally, when we tested a substrate that lacked TEG-biotin but possessed a 3′-P group (Substrate-3′P), we found that this substrate was ligated by CS1 with rates comparable to that of a biotinylated substrate (Figs. 4B, C). Therefore, the apparent dependence of CS1 on the substrate 3′-TEG-biotin was, in fact, a requirement for a 3′-phosphate group on the substrate. CS2, CS4, and CS5, like CS1, ligated to Substrate-3′P revealing the isolated ribozymes to represent a new class of ligases that catalyze a reaction between its 5′-triphosphorylated end and the 2′-OH group of a 3′-phosphorylated oligoribonucleotide (Fig. 4C, D).

**Figure 4.**
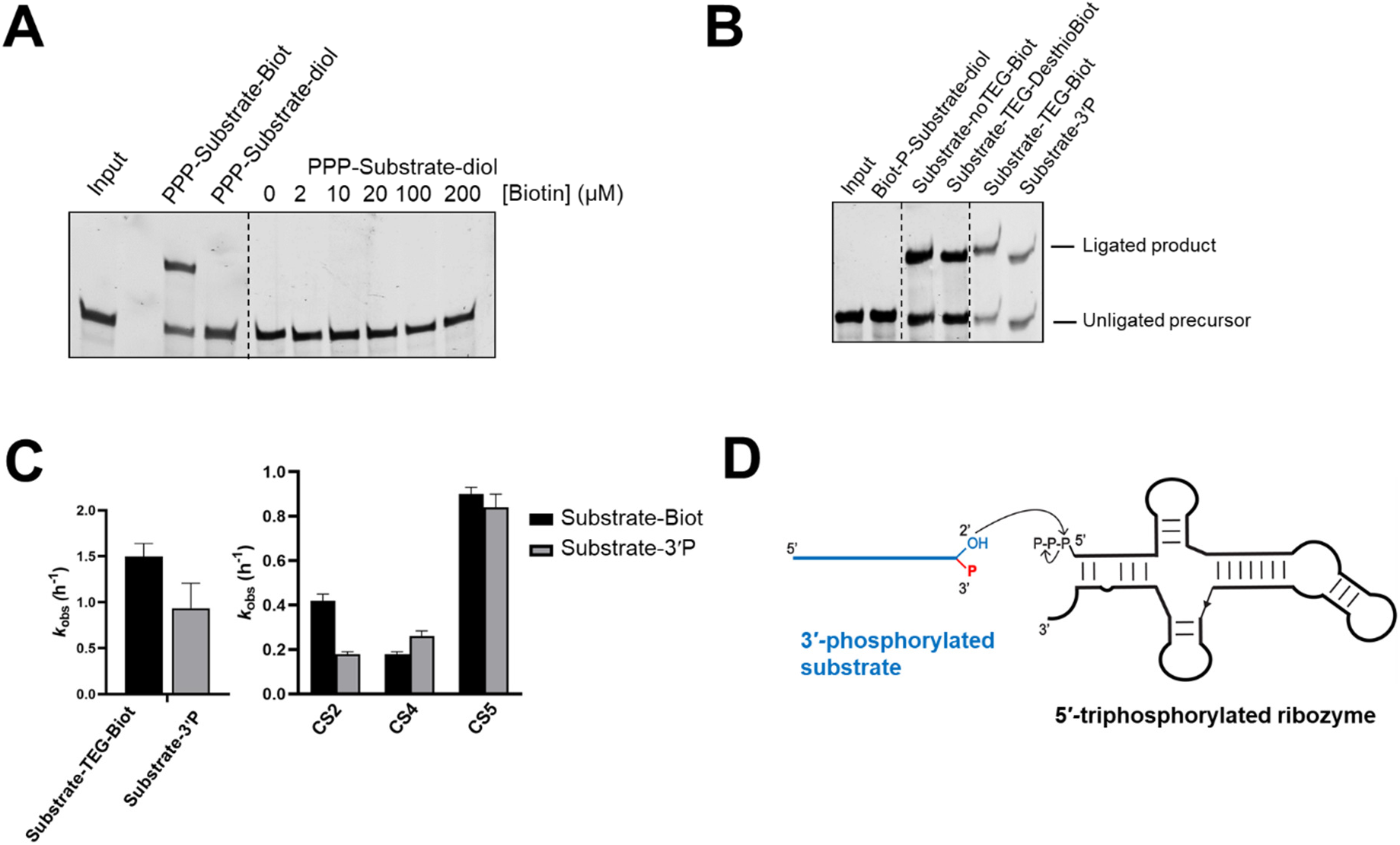
Ligation requires a 3′-phosphate group on the substrate. **A.** Ligation with an unbiotinylated substrate containing a terminal cis-diol is not rescued upon the addition of free biotin. **B.** Ligation is abolished when the TEG-biotin moiety is moved to 5′ end of the substrate, but a substrate with the biotin group replaced by a desthiobiotin group retains the ability to be ligated. A substrate with a 3′-phosphate group is active for ligation. **C.** Ligation rates for CS1, CS2, CS4, and CS5 with substrates containing terminal TEG-biotin or 3′-phosphate groups are comparable. **D.** The isolated ligase activity involves the nucleophilic attack of the 2′-OH group of a 3′-phosphorylated substrate on the 5′-triphosphate group of the ribozyme. The secondary structure of the ribozyme depicted here is the SHAPE-derived structure of CS1 (see Extended Data Fig. 15). Ligation reactions contained 1 µM ribozyme, 1.2 µM template (for CS2, CS4, and CS5) and 2 µM RNA substrate in 100 mM Tris-HCl (pH 8.0), 300 mM NaCl, and 100 mM MgCl_2_. Biotin was added to the reaction as indicated in (A).

As the isolated ribozymes catalyze substrate ligation to their 5′ end as opposed to their 3′ end, we were surprised that these sequences could be reverse transcribed at 42 °C using a primer (RT primer) that binds to its 3′ end by forming just 4 base-pairs. This indicates that it was possible to survive this step in the selection cycle by exploiting weak interactions between the RT primer and the ribozyme 3′ end (Extended Data Fig. 8). Because this ligation pathway joins the 2′ end of the substrate to the 5′ end of the ribozyme, the external template cannot be responsible for bringing these two ends into close proximity, as it does for the expected ligation junction between the 3′ end of the ribozyme and the 5′ end of the substrate (Extended Data Fig. 8). Furthermore, since the CS1 ribozyme does not require the external template, we were surprised to find that CS2, CS4, and CS5 did require an external template for activity (Extended Data Fig. 9A). We hypothesized that this template dependence was due to functional base-pairing interactions between the ribozyme 3′ sequence (‘primer’) and the template (Extended Data Fig. 9B). Disrupting base pairs between the primer and the template through template mutations eliminated ligation, which was rescued by compensatory mutations in the ribozyme 3′ primer, supporting this hypothesis (Extended Data Fig. 9C). It is possible that base-pairing interactions between the template and both the 3′ end of the ribozyme and the 5′ end of the substrate make this three-component reaction pseudo-intramolecular, thereby stimulating ligation by CS2, CS4, and CS5.

The essential role for the substrate 3′-phosphate is underscored by the rate acceleration of over 5 orders of magnitude relative to the background ligation rate (background ligation between a 5′-PPP-RNA oligonucleotide corresponding to the first 25 nt of the ribozymes and a 5′-FAM labeled 3′-phosphorylated substrate was measured as 2.4 x 10^-6^ h^-1^ at pH 8 and 100 mM Mg^2+^). Although a 3′-P can, in principle, accelerate substrate ligation by activating the vicinal 2′-OH nucleophile by acting as a general base catalyst, similar ligation rates with Substrate-3′P and Substrate-TEG-Biot discount this potential catalytic role. A more likely role of the 3′-P is in localizing catalytic divalent cations to the active site. Crystal structures of the L1 ligase and the class I ligase ribozymes, both of which catalyze ligation between the substrate 3′-OH and a ribozyme 5′-PPP, reveal that an active site Mg^2+^ interacts with a non-bridging oxygen on the α-phosphate of the ribozyme 5′-PPP and also with the ribozyme phosphodiester backbone.^17, 18^ In addition, the structure of the class I ligase suggests a possible interaction between Mg^2+^ and the 3′-O nucleophile.^25^ Similar interactions in these ligases between active site Mg^2+^ ions and the 3′-P group of the substrate and pyrophosphate leaving group of the ribozyme 5′-PPP could be important in catalysis. To examine the nature of the interaction between the terminal phosphate group and Mg^2+^, we measured reaction kinetics with substrates containing either a terminal phosphate or a thiophosphate in the presence or absence of the thiophilic cation Cd^2+^, in the background of Mg^2+^. We observed a 2-fold reduction in ligation rate with a terminal thiophosphate-containing substrate (Substrate-3′sP). When the reaction was supplemented with Cd^2+^ (25 µM or 1 mM), ligation was restored with a metal rescue value of ∼2.5 (Extended Data Fig. 10A, B). More pronounced thio effects and metal rescue values have been reported in ribozymes where Mg^2+^ directly interacts with either of the non-bridging oxygens of the phosphate group.^19^ A diminished effect is expected as our experiment does not involve a stereospecific O to S substitution in the phosphate. The modest effect we observed might also reflect a weak inner-sphere Mg^2+^ interaction. Substitution of Mg^2+^ with the divalent metal ions Ba^2+^, Ca^2+^, Co^2+^, Cu^2+^, Mn^2+^, Ni^2+^ abolished ligation, consistent with a possible catalytic role for Mg^2+^ (Extended Data Fig. 10C).

### Specific capture and amplification of 3′-phosphorylated RNA

RNAs with terminal phosphates in the form of 2′, 3′-cyclic phosphates (cP) and 3′-phosphates (3′-P) are generated as products of enzymatic cleavage pathways in RNA processing and maturation and have been implicated in diseases including cancers, amyotrophic lateral sclerosis, tuberculosis, and Parkinson′s disease.^20–25^ However, cleaved RNAs constitute a poorly characterized portion of the transcriptome, primarily because they remain invisible to standard library preparation protocols used in high-throughput sequencing. This is due to the inability of 3′-phosphorylated cleaved RNAs to ligate to the 3′ sequencing adapter because they lack free 3′-hydroxyl groups. The handful of recently reported methods for sequencing terminal phosphate-containing RNAs cannot distinguish between cP and 3′-P-containing RNAs due to the promiscuity of the RNA ligases used (*Arabidopsis thaliana* tRNA ligase or RtCB ligase) or rely on indirect enrichment of cP-RNAs via periodate cleavage of RNA terminal diols.^26–29^. As the ribozymes identified in this work show absolute discrimination between 3′-OH and 3′-P RNA termini we explored their potential application as reagents for the specific enrichment of 3′-phosphorylated RNA from total cellular RNA.

We found the CS1 has a low *K*_M_ value of 0.1649 ± 0.042 µM and a high catalytic efficiency (*k*_cat_/ *K*_M_ value) of 27.65 µM^-1^ h^-1^ (7670 M^-1^ s^-1^) indicating that CS1 could function as a potential reagent for enriching 3′-P-RNAs (Extended Data Fig. 11). We spiked in different concentrations a 5′ FAM-labeled 3′-phosphorylated substrate (FAM-Target RNA-3′P) into an *E. coli*-derived tRNA mix and incubated this mixture with CS1. The appearance of a band corresponding to the ligated product even when tRNAs were in 60-fold excess supports the ribozyme’s ability to enrich 3′-phosphorylated RNAs from a heterogeneous mixture of cellular RNA (Fig. 5A). The fluorescence signal from the ligated product increased linearly with substrate concentration between 0.05 uM to 0.4 uM, suggesting a potential for quantitative detection of 3′-P-RNAs (Fig. 5B). The captured 3′-P-RNAs must be first reverse transcribed and then PCR-amplified to generate material that can be sequenced (Extended Data Fig. 12A). To see whether the 3′-P next to a 2′-5′ phosphodiester linkage would prevent reverse transcription, we compared RT-PCR before and after removing the phosphate group by shrimp alkaline phosphatase (SAP). Interestingly, the target RNA was amplified even without SAP treatment, showing that reverse transcriptase can copy across the unusual 2′-5′ phosphodiester linkage harboring an adjacent 3′-phosphate (Extended Data Fig. 12B).

**Figure 5.**
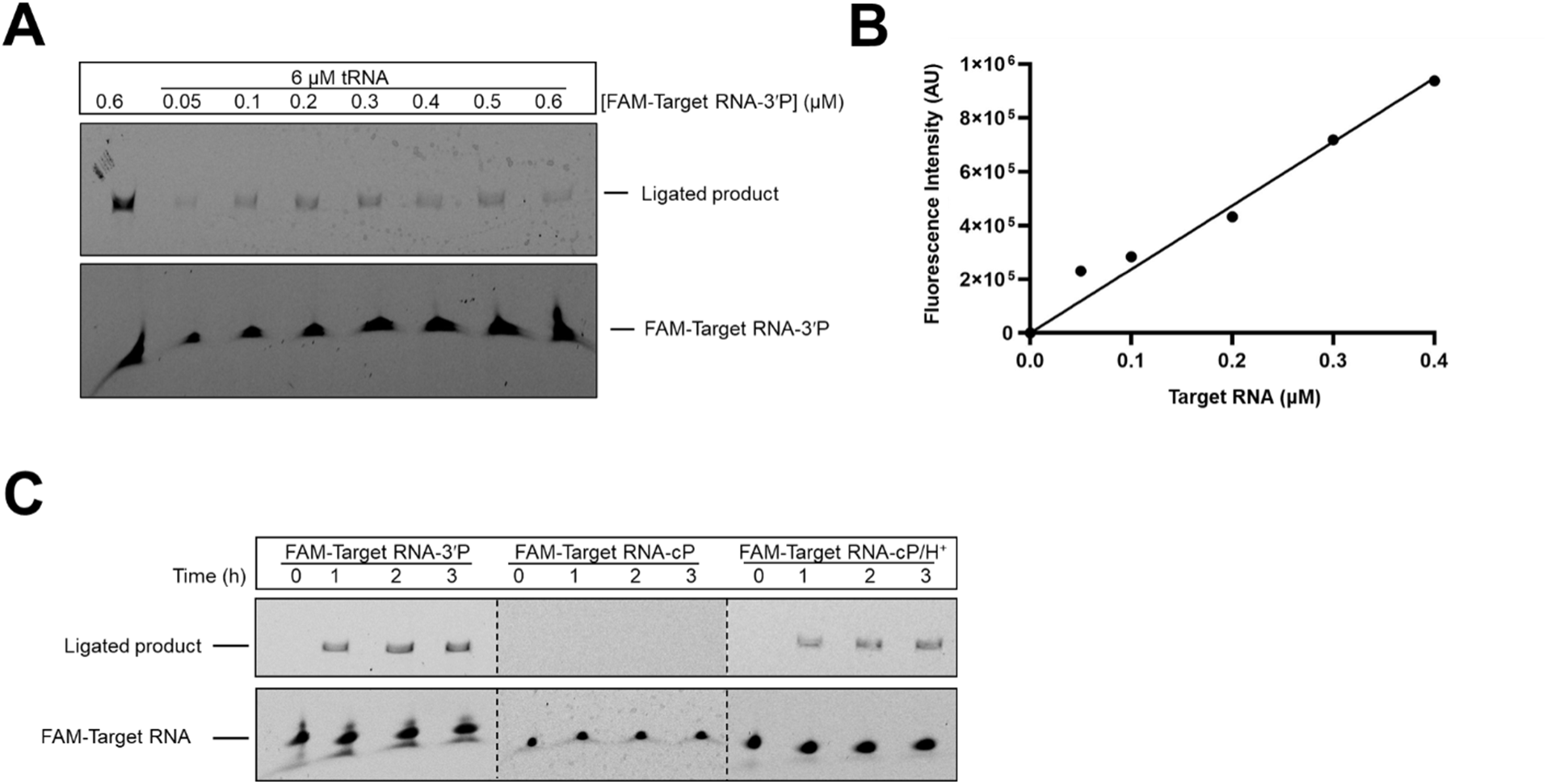
Ribozyme-assisted capture of 3′-phosphorylated RNA. **A.** CS1 captures a FAM-labeled 3′-phosphorylated substrate, FAM-Target RNA-3′P, from a heterogeneous mixture of cellular tRNAs. **B.** Capture, as measured by the fluorescence intensity of the ligated product, is linear in response to substrate concentration between 0.05 uM and 0.4 uM. **C.** CS1 shows specificity toward substrate 3′ termini. It specifically ligates to substrates with 3′-phosphate groups, while being inert to those terminating in 2′, 3′-cyclic phosphate groups. Ligation reactions contained 1 µM ribozyme, CS1, and the indicated amounts of FAM-Target RNA-3′P (A, B) or 2 µM RNA targets (FAM)Target RNA-3′P or FAM-Target RNA-cP) in 100 mM Tris-HCl (pH 8.0), 300 mM NaCl, and 100 mM MgCl_2_.

These ribozymes may also be used to indirectly capture cP-RNAs by converting their cP ends to 3′-P (and 2′-P) on mild acidification. Upon incubating a cP-containing substrate (FAM-Target RNA-cP) pre-treated with 10 mM HCl (0 °C/3 h)^29^ with CS1, we observed a steady increase in ligation with increasing incubation time, indicating cP-RNA capture. FAM-Target RNA-cP, not subjected to the acidification step, was not captured by the ribozyme (Fig. 5C). As self-cleaving ribozymes generate cleavage products with cP ends, this ribozyme-assisted enrichment method, in conjunction with standard RNA-seq, may be used in high-throughput screens to discover new self-cleaving ribozymes.^30, 31^

For maximal utility as RNA sequencing reagents, ribozymes must enrich RNAs in a sequence-general manner. Therefore, we tested the substrate scope of ribozyme-assisted target capture. We tested CS1 against truncated versions of the Substrate-3′P, which has the sequence: 5′-ACCACCGCAUUCCGC**A**p-3′, where **A** contains the 2′-OH nucleophile. CS1 captured a target representing the last 8 residues of Substrate-3′P (5′-AUUCCGC**A**p-3′) but was unable to ligate an oligoribonucleotide representing its first 8 residues (5′-ACCACCGCp-3′) (Extended Data Fig. 13A). Target 5′-AUUCCGC**A**p-3′ was further truncated to two 5 nt pieces and tested for ligation. Once again, the piece containing the nucleophilic adenine (5′-CCGC**A**p-3′) was captured but 5′-AUUCCp-3′ was not (Extended Data Fig. 13A). Shorter oligomers were not tested due to the difficulty in resolving captured products from the ribozyme by denaturing gel electrophoresis. Substrates where their terminal A was replaced by C or U could not be captured by CS1, and a substrate with a 3′ terminal G showed ∼500-fold slower ligation with CS1 (Extended Data Fig. 13B, C). CS2, CS4, and CS5 showed similar dependence on the identity of the 3′ terminal residue of the substrate (Extended Data Fig. 14). These results indicate the importance of the adenine nucleotide at the 3′ end of the substrate. Similarly, the ribozyme 5′ end was constrained to a guanine; a G to A mutation in all four ribozymes abolished ligation with all four substrate variants (Extended Data Figs. 13, 14). These constraints on both ligation junction nucleotides suggest that there is an internal template that brings them together to allow ligation, however the identity of this internal template, if it exists, remains to be defined. The loss of activity in the CS1_5’A mutant is consistent with a G1-C74 base pair in stem P1 as suggested by the SHAPE-derived secondary structure of CS1 (Extended Data Fig. 15). Base pairing at the ribozyme 5′ end may rigidify the ligation junction, thereby facilitating reactive interactions with the substrate. The unpaired 5′-UGU-3′ present close to the putative ligation junction (nucleotides 77-79) may base pair with the 3′ terminal 5′-GCA-3′ portion of the substrate facilitating substrate binding and positioning. This is consistent with the compromised CS1-catalyzed ligation observed with substrates possessing a non-adenine residue at its 3′ end.

## DISCUSSION

Directed evolution of RNA has made it possible to experimentally access RNA sequences with new functions from existing ones. Although in most cases, RNAs with a desired target function have been identified, there are several examples where unexpected functions have emerged.^4, 5, 8^ Thus a single selection condition may give rise to multiple RNA activities as long as the activity allows the sequences to survive the selection pressure, i.e., exhibit the required fitness. In this work, we isolated ligase ribozymes that catalyze a reaction between the 2′-OH group of an RNA substrate and the 5′-PPP group of the ribozyme. The isolated activity is the opposite of the targeted activity – the nucleophilic attack of the ribozyme 3′-OH on the substrate 5′-PPP group. While ribozymes with 2′-5′ ligase activity have been identified before,^6, 32^ the ligases identified in this work present a new variation with a strict requirement for a 3′-phosphate, vicinal to the nucleophilic 2′-OH group on the substrate. The similarities of this ribozyme ligase reactivity to protein-based RNA repair ligases such as tRNA ligase and RtcB ligase ^9, 20^ are worth noting. tRNA ligase catalyzes a reaction between the 5′-adenyl-pyrophosphate (App) moiety of one RNA substrate and a 3′-OH group of the other, where the second RNA substrate contains a 2′-phosphate vicinal to the nucleophilic 3′-OH. This pathway resembles the ligase ribozymes reported here, wherein ligation involves a 5′-triphosphate of one RNA and the terminal 2′-OH of the other RNA, in the presence of a vicinal 3′-phosphate. RtcB ligase uses 3′-P-RNAs as substrates; however, there, the 3′-phosphate participates directly in reaction, unlike the ligase ribozymes we isolated.

It is important to recognize that designing a selection strategy to intentionally isolate 3′-P-specific ligases where the 3′-phosphate group does not directly participate in the ligation reaction would be extremely challenging. However, we unintentionally isolated four prominent classes of ribozyme ligases with this activity. These ligases diverge considerable from the parental AI-Ligase (by 10-16 mutations) and from each other (by 11-29 mutations). The predominance of 3′-P-specific ligases isolated from our selection suggests that our selection scheme, with minor modification, may be used to isolate more efficient catalysts, including those with expanded substrate scopes. Optimized ribozymes that ligate to RNA substrates regardless of the identity of their terminal residue, unlike those reported here, have potential to act as reagents for the transcriptome-wide probing of terminal phosphate-containing RNA cleavage products.^20^ The ability to generate a complete catalog of cellular cP- and 3′-P-RNAs under various cellular conditions and in health and disease will expand our understanding of RNA cleavage and repair pathways and potentially identify disease biomarkers.

Going back in time, RNAs containing terminal phosphates would have been generated on the early Earth by nonspecific phosphorylation at their 5′ and 3′ termini, and due to cleavage of the RNA backbone followed by hydrolysis of the 2′-3′-cyclic phosphate.^33–35^ Excessive RNA cleavage would prevent efficient genome replication in early RNA-based lifeforms. Ribozymes capable of ligating cleaved RNAs to regenerate a continuous RNA backbone provide a salvage pathway by which the information contained within the cleaved RNA may be preserved (Extended Data Fig. 16). 5′ RNA cleavage products containing terminal 2′,3′-cyclic phosphates are easily acid-hydrolyzed to 3′-phosphates (and 2′-phosphates) generating substrates for the ribozymes reported in this work.^29^ 3′ cleavage products containing 5′-hydroxyl groups may be triphosphorylated by ribozymes in the presence of cyclic-trimetaphosphate, an abundant prebiotic reagent.^36–38^ Ligation between 5′-triphosphorylated RNAs and 3′-phosphorylated RNAs catalyzed by trans versions of these ribozymes would restore the original RNA sequence before cleavage, with the 2′-5′ phosphodiester linkage being the only point of distinction. The presence of a 2′-5′ phosphodiester linkage in the ligated product does not prevent its replication^39^ and RNAs containing interspersed 2′-5′ linkages have been shown to retain their functions.^40^ The viability of prebiotically plausible backbone repair pathways that convert internal 2′-5′ linkages to 3′-5′ linkages^41^ support this proposed ribozyme-assisted salvage pathway. This pathway complements a recently proposed nonenzymatic pathway for ‘patching’ cleaved RNAs containing 3′-phosphates via the formation of 3′-5′ pyrophosphate linkages.^42^ In the wider context of the origin and evolution of early life, our results demonstrate the ease with which diverse RNA catalysts can emerge under the same selection pressure, further bolstering the case for the existence of a primordial RNA-based biology on the early Earth.

## METHODS

### Materials

All chemicals were purchased from Sigma, unless otherwise specified. The hydrochloride salts of 1-ethyl-3-(3-dimethylaminopropyl) carbodiimide and 2-aminoimidazole were purchased from Alfa Aesar and Combi-Blocks, Inc, respectively. Enzymes were purchased from New England Biolabs unless mentioned. SYBR Gold Nucleic Acid Gel Stain was purchased from ThermoFisher Scientific. 100% ethanol was purchased from Decon Laboratories, Inc. QIAquick PCR purification kits were purchased from Qiagen. The Sequagel-UreaGel concentrate and diluent system for denaturing polyacrylamide gels was purchased from National Diagnostics. Oligonucleotides were purchased from Integrated DNA Technologies (IDT) except for PPP-Substrate and PPP-Substrate-Biot, which were purchased from Chemgenes.

### RNA preparation and substrate activation

Ribozyme constructs were prepared by in vitro transcription of dsDNA templates generated by PCR of single-stranded DNA. These dsDNA templates contained 2′-*O*-methyl modifications on the last two nucleotides of the template strand to reduce transcriptional heterogeneity at the 3′ end of the RNA.^43^ Each transcription reaction containing 40 mM Tris-HCl (pH 8), 2 mM spermidine, 10 mM NaCl, 25 mM MgCl_2_, 10 mM dithiothreitol (DTT), 30 U/mL RNase inhibitor murine, 2.5 U/mL thermostable inorganic pyrophosphatase (TIPPase), 4 mM of each NTP, 30 pmol/mL DNA template, and 1U/µL T7 RNA Polymerase were incubated at 37°C for 2-3 h. The dsDNA template was removed by DNase I-digestion (5U/mL at 37° C for 30 min) and the RNA was extracted with phenol-chloroform-isoamyl alcohol (Invitrogen), ethanol precipitated, and purified by 10% denaturing PAGE.

CS1 variants with 5′-phosphate (5′-P) and 5′-hydroxyl (5′-OH) groups were generated by the splinted ligation of three RNA oligonucleotide pieces (Supplementary Table 1). The first piece contained either a 5′-P or a 5′-OH modification. The second and third pieces were 5′-monophosphorylated to enable ligation by T4 RNA ligase 2. 60 pmol of each piece were incubated with 40 pmol two DNA splints each at 90 °C for 3 min, followed by a 10 min incubation at 30 °C, which was followed by the addition of 1U/ µL RNA ligase 2 and 1X T4 RNA ligase buffer. The 20 µL reaction was incubated at 30 °C for 2 h, the RNA was extracted with phenol-chloroform-isoamyl alcohol, and purified by 10% denaturing PAGE.

The 5′-monophosphorylated oligonucleotide corresponding to the AIP-Substrate (Supplementary Table 1) was activated by as previously described.^11^ Briefly, it was reacted with 0.2 M 1-ethyl-3-(3 dimethylaminopropyl) carbodiimide (HCl salt) and 0.6 M 2-aminoimidazole (HCl salt, pH adjusted to 6) in aqueous solution for 2 h at room temperature. Salts were removed by four or five washes (200 µL water per wash) in 3kDa cutoff Amicon Ultra spin columns. This was followed by reverse phase analytical HPLC purification using a gradient of 98% to 75% 20 mM TEAB (triethylamine bicarbonate, pH 8) versus acetonitrile over 40 min.

### Ribozyme-catalyzed RNA ligation assays

Ligation reactions contained 1 µM ribozyme, 1.2 µM RNA template, and 2 µM RNA substrate in 100 mM Tris-HCl pH 8.0, 300 mM NaCl, and either 100 mM MgCl_2_, unless specified. Aliquots were quenched with 5 volumes of quench buffer (8 M urea, 100 mM Tris-Cl, 100 mM boric acid, 100 mM EDTA) and analyzed on a 10% denaturing PAGE. Gels were stained using SYBR Gold and imaged on an Amersham Typhoon RGB instrument (Cytiva), Scans were analyzed in ImageQuant IQTL 8.1. Kinetic data were nonlinearly fitted to the modified first order rate equation, y = A (1 – e-*^k^*^x^), where A represents the fraction of active complex, *k* is the first order rate constant, x is time, and y is the fraction of ligated product in GraphPad Prism 9.

### Confirmation of a 2′-5′ phosphodiester bond in the ligated product

#### Terminator 5′-3′ exonuclease stop assay

∼6 pmol of the purified ligated products obtained from an overnight ligation reaction between CS1 and PPP-Substrate-Biot or P-Substrate-Biot were digested with 2 U of Terminator (Lucigen) in a 20 μL reaction in the presence of 1X Terminator buffer A at 30 °C for 2 h. The reaction was cleaned up using Zymo RNA Clean & Concentrator-5 kit (Zymo Research) and the RNA was eluted in 6 μL milliQ water. 10 μL of quench buffer (8 M urea, 100 mM Tris-Cl, 100 mM boric acid, 100 mM EDTA) was added to it, followed by 10% denaturing PAGE.

#### RNase R 3′-5′ exonuclease stop assay

∼5 pmol of the purified ligated product obtained from an overnight ligation reaction between CS1 and Substrate-Biot was digested with 20 U of RNase R (Lucigen) in a 10 μL reaction in the presence of 1X RNase R buffer at 50 °C for 1 h. The reaction was quenched by adding 0.3 μL of 0.5 M EDTA and the enzyme was heat-inactivated at 95 °C for 3 min. 10 μL of quench buffer (8 M urea, 100 mM Tris-Cl, 100 mM boric acid, 100 mM EDTA) was added to the reaction, followed by 20% denaturing PAGE.

### Secondary structure determination by SHAPE probing

SHAPE probing of CS1 was performed using a protocol reported in Walton *et al.*, 2020.^2^ Briefly, 100 pmol of the RNA construct containing 5′ and 3′ SHAPE cassettes (CS1_SHAPE; see Supplementary Table 1) was folded in 100 mM Tris-HCl, pH 8, 250 mM NaCl, 10 mM Mg^2+^ and divided into ‘modification’ and ‘control’ tubes. ∼40 mM of the SHAPE reagent, 1M7 (dissolved in DMSO), was added to the ‘modification’ tube and equal volumes of DMSO was added to the ‘control’ tube. Both modified and unmodified RNAs were reverse transcribed using 40 pmol of a 5′ FAM-labeled primer (SHAPE_RT_primer; Supplementary Table 1) and Superscript III reverse transcriptase (Invitrogen) (Supplementary Table 1). Reverse transcription of 30 pmol unmodified RNA in the presence of each of the four ddNTPs and 25 pmol SHAPE_RT_primer was used to generate sequencing lanes. Quenched reactions were analyzed by 10% denaturing PAGE. Normalized SHAPE reactivities were calculated by first excluding the most reactive nucleotide position and then dividing reactivities at each position by the average of the 10% most reactive positions. Normalized SHAPE reactivities were used to constrain secondary structure prediction in the RNAstructure program.^44^

### Capture and amplification of 3′-phosphorylated RNA

To demonstrate the capture of 3′-phosphorylated RNA from a mixture of cellular RNA, 0.3 μM CS1 was added to 6 μM *E. coli* tRNA mix (Millipore Sigma) that was spiked with different concentrations (0.1, 0.2, 0.3, 0.4, 0.5, 0.6 μM) of 5′ FAM-labeled RNA target (FAM-Target RNA-3′P) under standard ligation conditions. Aliquots were taken after 3 h, quenched with 9 μL of quench buffer and analyzed by 10% denaturing PAGE.

To demonstrate the capture of target RNA containing 2′-3′-cyclic phosphate, first the 2′-3′-cyclic phosphate-containing RNA (FAM-Target RNA-cP) was generated via hairpin ribozyme-mediated cleavage of FAM-HP_sub (Supplementary Table 1). 4 μM hairpin ribozyme, HP_ribozyme, was heated at 70 °C for 1 min followed by incubation at room temperate for 5 min. The solution was incubated at 37 °C for 15 min after adding Tris-HCl (pH 8) to a final concentration of 300 mM. This was followed by the addition of 100 mM MgCl_2_ and 2 μM FAM-HP_sub. The cleavage reaction was incubated at 37 °C for 5 h. The 5′ cleaved product containing 2′, 3′-cyclic phosphate (FAM-Target RNA-cP) was purified by 10% denaturing PAGE. Purified FAM-Target RNA-cP (42 pmol) was converted to FAM-Target RNA-3′P by adding 10 mM HCl and incubating for 3 h on ice. The RNA was then ethanol precipitated. The target capture reaction contained 2.5 pmol FAM-Target RNA-cP and 6 pmol CS1 under regular ligation conditions and was analyzed by 10% denaturing PAGE.

To demonstrate the efficient amplification of captured 3′-phosphorylated RNA, the ligated product obtained from a reaction between CS1 and Substrate-3′P was incubated with 10 U shrimp alkaline phosphatase (SAP; New England Biolabs) at 37 °C for 30 min, followed by heat inactivation at 65 °C for 20 min. The inactivated SAP was removed by Zymo RNA Clean & Concentrator-5 kit. The SAP-treated ligated product was reversed transcribed in a 20 μL reaction containing 10 μM RT primer, 2 mM dNTP mix, 2 mM DTT, 1X Protoscript buffer, and 5 U/μL Protoscript II reverse transcriptase, incubated at 37 °C for 3 h. Ligated product that was not treated with SAP was also reverse transcribed as a control. The reverse transcriptase was heat inactivated at 80 °C for 5 min and the cDNAs were purified with QIAquick PCR purification kit. The purified product was directly used as input for a PCR with primers, PCR_LigFwd_primer and PCR_Rvs_primer (Supplementary Table 1). The PCR cycle was: 1) 95 °C for 2 min; 2) 16 cycles of 94 °C for 30 s, 54 °C for 1 min., and 72 °C for 1 min.; 3) 72 °C for 10 min. The PCR product was purified using QIAquick PCR purification kit and analyzed by 10% denaturing PAGE.

## ACKNOWLEDGMENTS

We thank members of the Szostak lab, especially, Dr. Victor Lelyveld for valuable feedback on the project and members of the DasGupta lab for helpful comments on the manuscript. We thank Dr. Joseph Piccirilli for helpful discussions during the project. We also thank the MGH’s NextGen Sequencing Core for help with Illumina sequencing. J.W.S. is an Investigator of the Howard Hughes Medical Institute. This work was supported in part by a grant from the Simons Foundation (290363) to J.W.S. and University of Notre Dame Startup Funds to S.D.

## AUTHOR INFORMATION

### Other Authors

**Annyesha Biswas −** Department of Chemistry and Biochemistry, University of Notre Dame, Notre Dame, Indiana, 46556, USA.

**Zoe Weiss −** Harvard/Massachusetts Institute of Technology MD-PhD Program, Harvard Medical School, Boston, MA 02115

### Author Contributions

A.B., J.W.S., and S.D. designed research; A.B. Z.W., and S.D. performed research; Z.W. contributed new analytical tools, A.B., Z.W., J.W.S., and S.D. analyzed data; S.D., J.W.S. wrote the paper.

## ETHICS DECLARATIONS

Competing Interest Statement: The authors declare no conflict of interest.

## EXTENDED DATA

**Extended Data Fig. 1.**
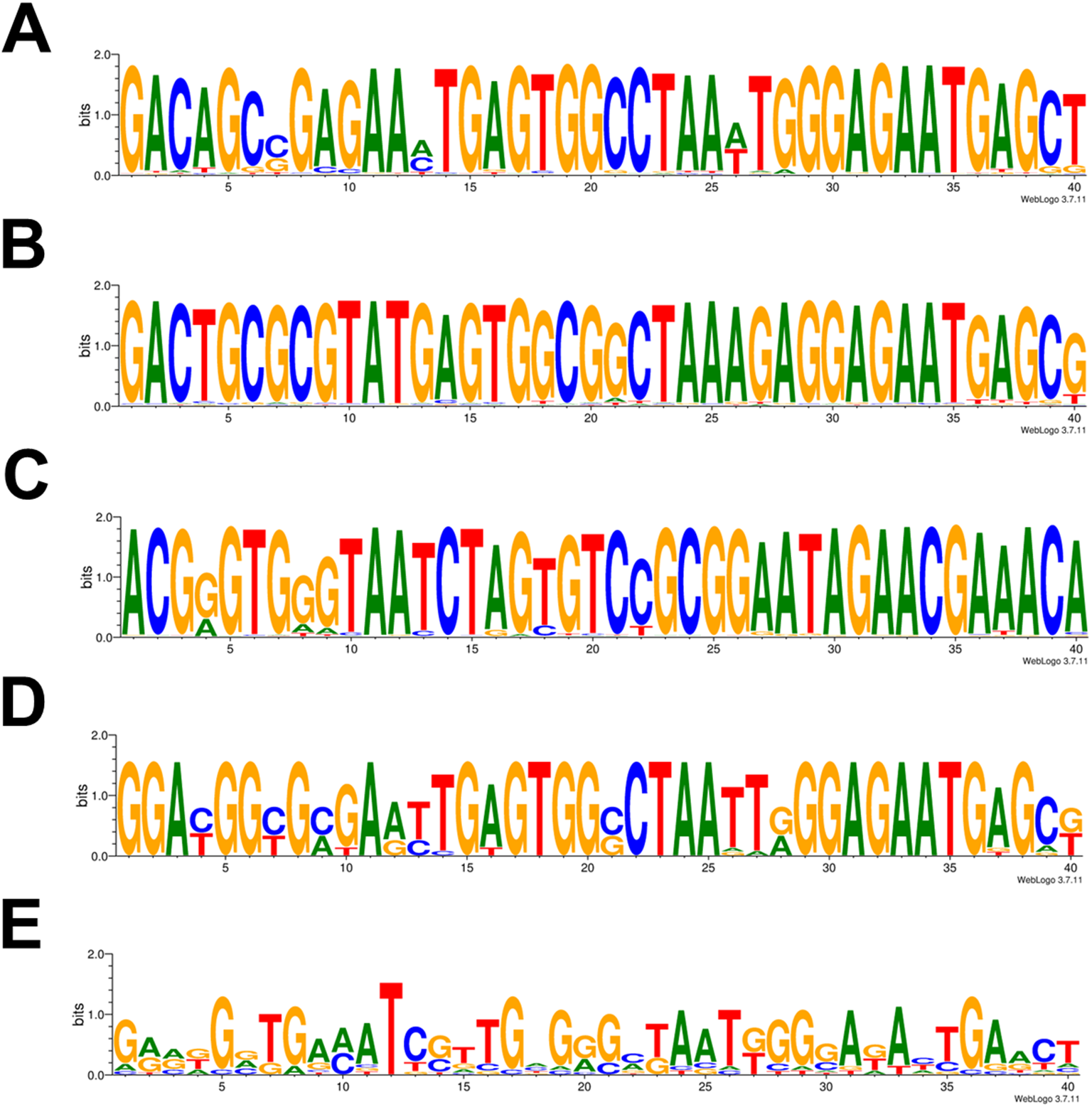
Sequence logos of the variable region in clusters 1-5. (A-E). The sequence logos show the consensus sequence for the 40 nt variable region and depicts the relative abundances of each nucleotide for each position. The sequence logo was generated by the WebLogo online server (https://weblogo.berkeley.edu/logo.cgi).

**Extended Data Fig. 2.**
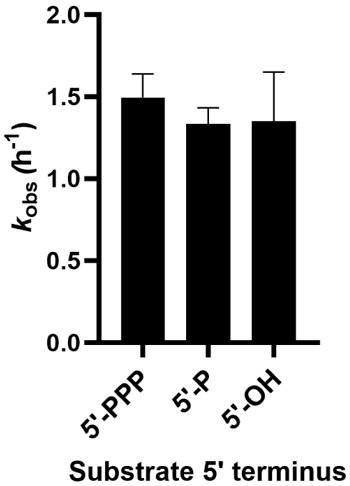
Ligation reactions with substrates containing 5′-triphosphate (5′-PPP), 5′-monophosphate (5′-P), or 5′-hydroxyl (5′-OH) groups are comparable. Ligation rates exhibited by CS1, the most abundant ligase isolated from the selection, are similar with substrates containing 5′PPP, 5′P, or 5′OH groups. Ligation reactions contained 1 µM ribozyme CS1 and 2 µM Substrate-Biot in 100 mM Tris-HCl (pH 8.0), 300 mM NaCl, and 100 mM MgCl_2_.

**Extended Data Fig. 3.**
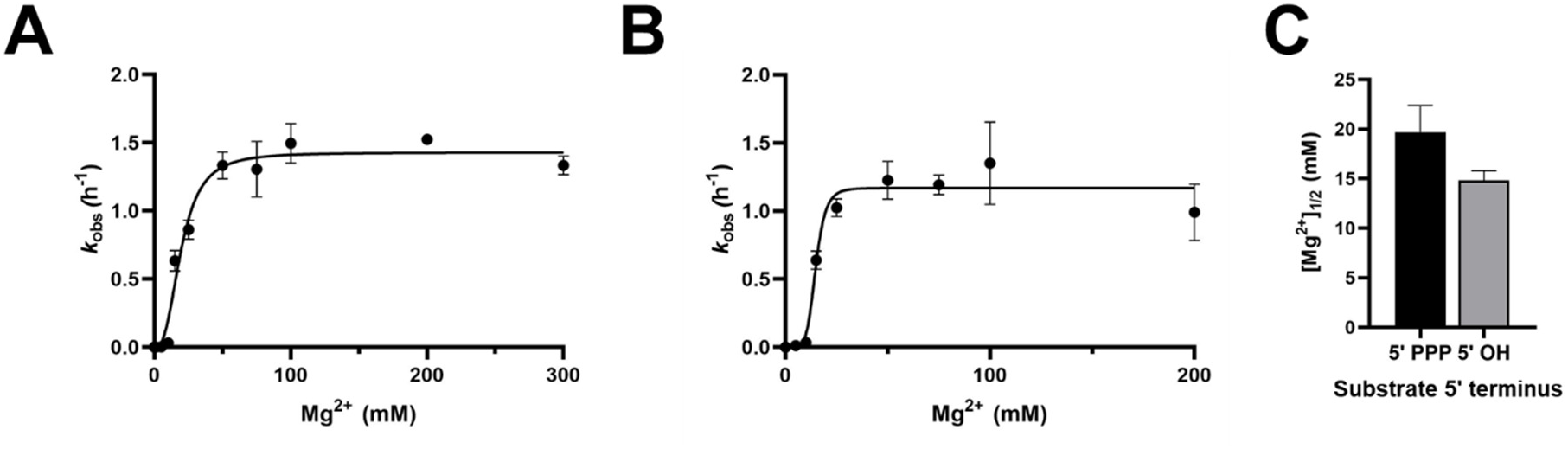
The effect of Mg^2+^ concentration on the rates of CS1-catalyzed ligation of substrates containing 5′-triphosphate (5′-PPP) and 5′-hydroxyl (5′-OH) groups is comparable. A. *k*_obs_ values with a PPP-Substrate-Biot increase steeply till [Mg^2+^] reaches 50 mM, and plateaus at higher concentrations. B. *k*_obs_ values with HO-Substrate-Biot increase steeply till [Mg^2+^] reaches 25 mM and plateaus at higher concentrations. **C**. Mg^2+^ titration data are plotted to the Hill Equation and yield [Mg^2+^]_1/2_ values of 19.7 ± 2.70 mM and 14.8 ± 2.98 mM for PPP-Substrate-Biot and HO-Substrate-Biot. Ligation reactions contained 1 μM ribozyme and 2 μM PPP-Substrate-Biot or HO-Substrate-Biot in 100 mM Tris-HCl (pH 8.0), 300 mM NaCl, and the indicated amounts of MgCl_2_.

**Extended Data Fig. 4.**
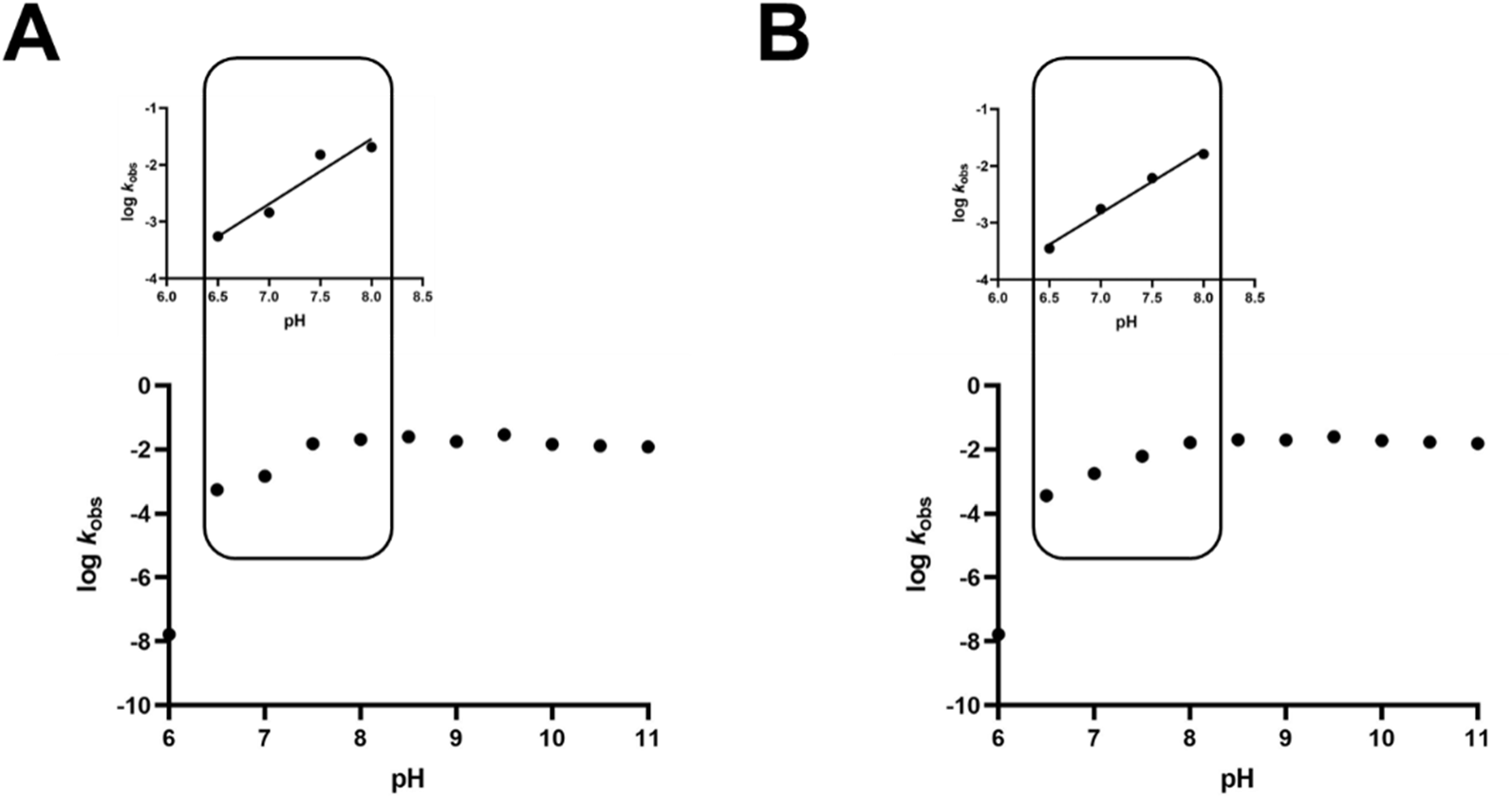
Comparable effect of pH on the rates of CS1-catalyzed ligation of substrates containing 5′-monophosphate (5′-P) and 5′-hydroxyl (5′-OH) groups. Log *k*_obs_ values increase to pH 8 and then remain approximately constant. The inset shows that the increase in ligation rate is log-linear in the range pH 6.5-8 with a slope of ∼1 with both P-Substrate-Biot and HO-Substrate-Biot. Ligation reactions contained 1 μM ribozyme and 2 μM P-Substrate-Biot or HO-Substrate-Biot, 300 mM NaCl, and 100 mM MgCl_2_ at the indicated pH values. The following buffers were used: MES: pH 6.0, 6.5; Tris: pH 7.0, 7.5, 8.0, 8.5, 9.0; Bis-tris propane: pH 9.5; CAPS: 10, 10.5, 11.

**Extended Data Fig. 5.**
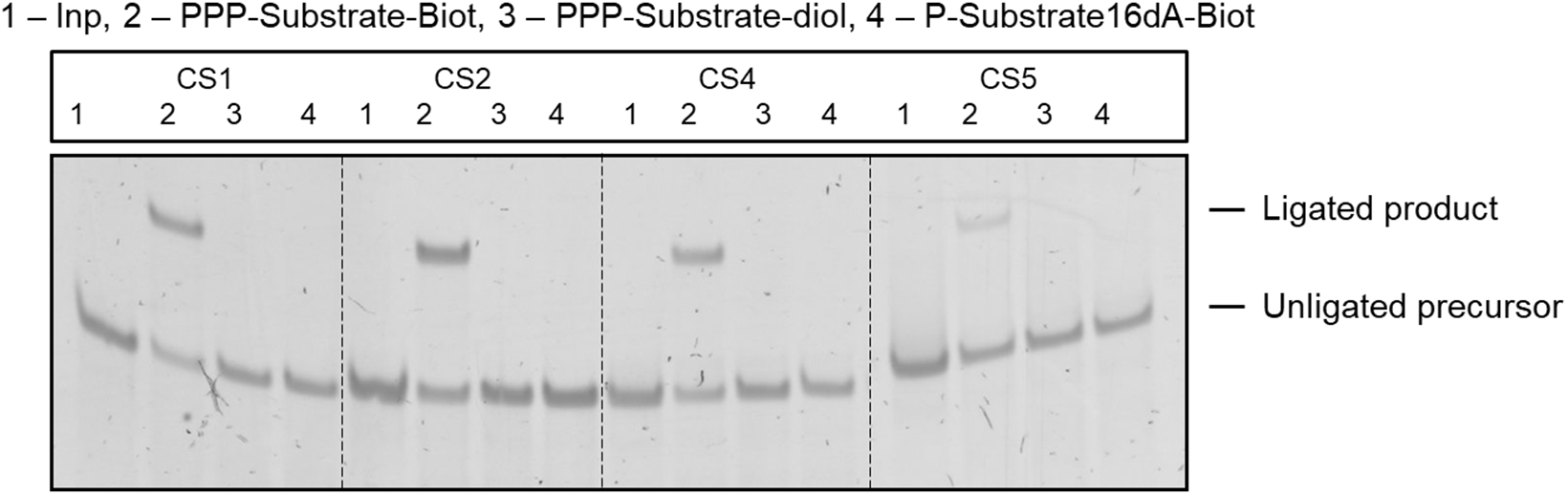
The reactivity of ribozymes toward substrates with different 3′ termini. CS1, CS2, CS4, and CS5 react with substrates that possess a 3′-TEG-biotin group but are inert toward substrates containing a terminal diol or terminating in a deoxynucleotide (dA). Lane 1 (Inp) is the input lane that contains only ribozyme. Ligation reactions contained 1 µM ribozyme and 2 µM RNA substrate in 100 mM Tris-HCl (pH 8.0), 300 mM NaCl, and 100 mM MgCl_2_.

**Extended Data Fig. 6.**
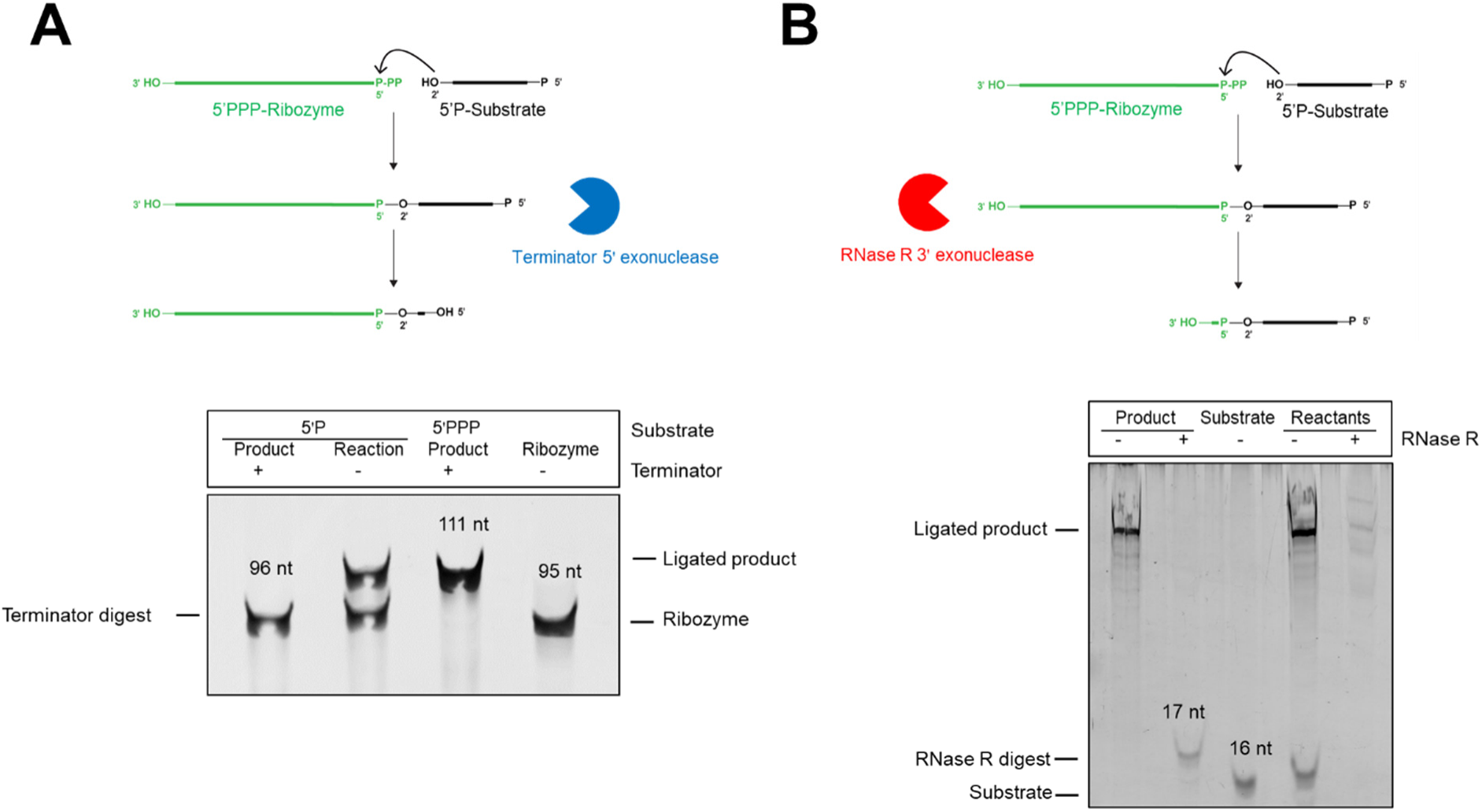
Ligation creates a 2 ′-5′ phosphodiester bond between the substrate and the ribozyme. The lengths of the partially digested products from separate 5′à3′ and 3′à5′ exonuclease digestions support the presence of a noncanonical 2′-5′. Between the substrate and the ribozyme **A.** The ligated product from the reaction between CS1 and P-Substrate-Biot is partially degraded by the 5′à3′ exonuclease, Terminator, to generate a digested product that is 1 nt longer than the ribozyme. The corresponding ligated product with PPP-Substrate-Biot is unreactive to Terminator degradation, serving as a negative control. **B.** The ligated product from the reaction between CS1 and Substrate-Biot is partially degraded by the 3′à5′ exonuclease, RNase R, to generate a digested product that is 1 nt longer than the substrate. A mixture of CS1 and the substrate, on the other hand, is completely degraded by RNase R. Exonuclease digestions were carried out according to vendor specifications (See ‘Confirmation of a 2′-5′ phosphodiester bond in the ligated product’ in Methods).

**Extended Data Fig. 7.**
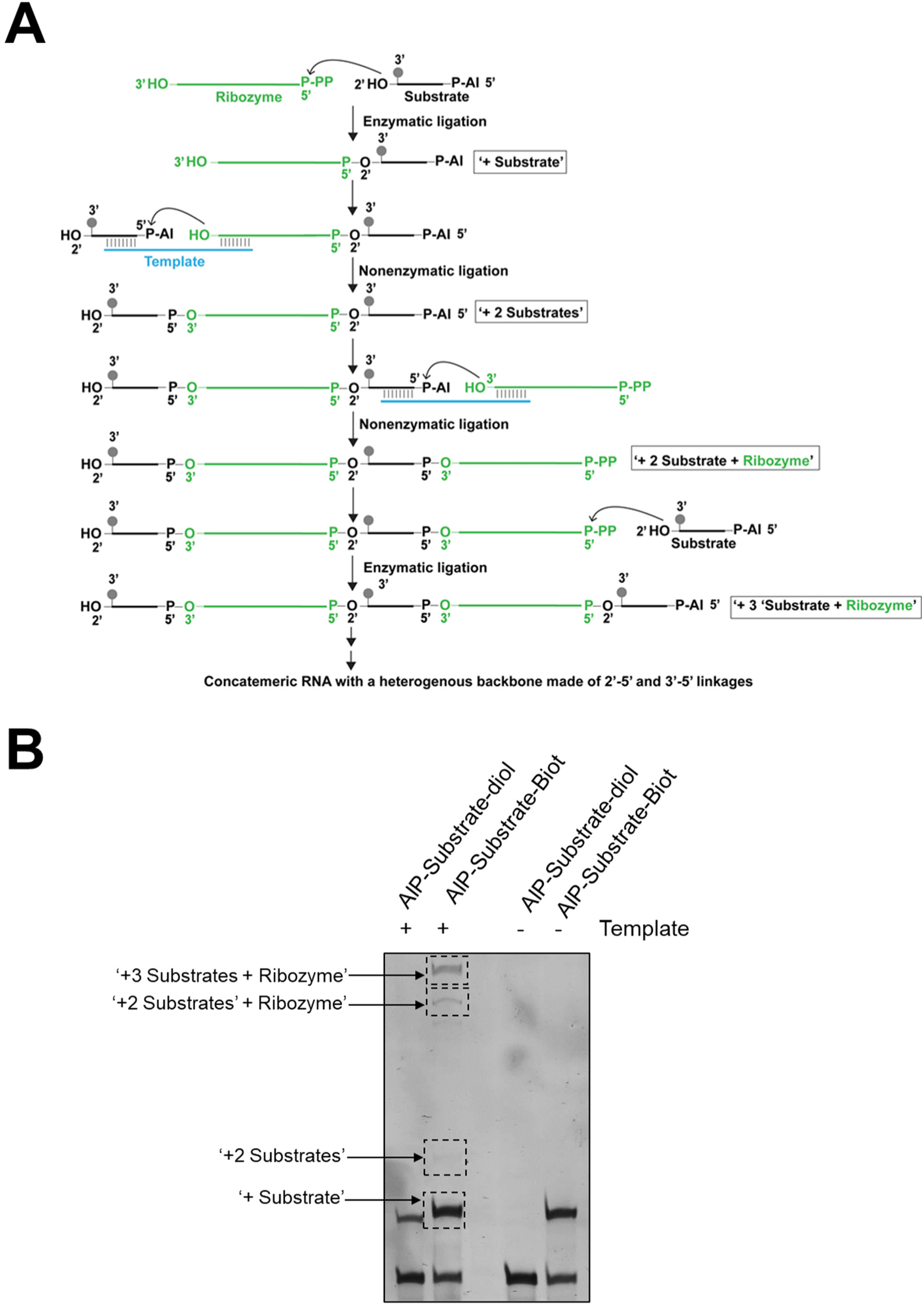
Ligation with 5′ 2-aminoimidazole-activated, 3′ biotinylated substrates produce concatemeric RNA containing both 2′-5′ and 3′-5′ linkages. A. A combination of ribozyme-catalyzed and nonenzymatic ligation generates concatemeric RNA. B. Higher-order bands corresponding to concatemeric ligation products are observed when CS1 is incubated with AIP-Substrate-Biot in the presence of a template at 100 mM Mg^2+^. These bands are not observed in the absence of a template because detectable nonenzymatic ligation required a template. Using an AIP substrate without TEG-biotin also fails to generate higher-order products because the ribozyme does not ligate RNAs with terminal diols. The ‘+substrate’ band observed with AIP-Substrate-diol in the presence of a template is due to nonenzymatic ligation. The gray circle represents the ‘TEG-Biotin’ moiety. Ligation reactions contained 1 μM ribozyme, 1.2 μM RNA template, and 2 μM RNA substrate in 100 mM Tris-HCl (pH 8.0), 300 mM NaCl, and 100 mM MgCl_2_.

**Extended Data Fig. 8.**
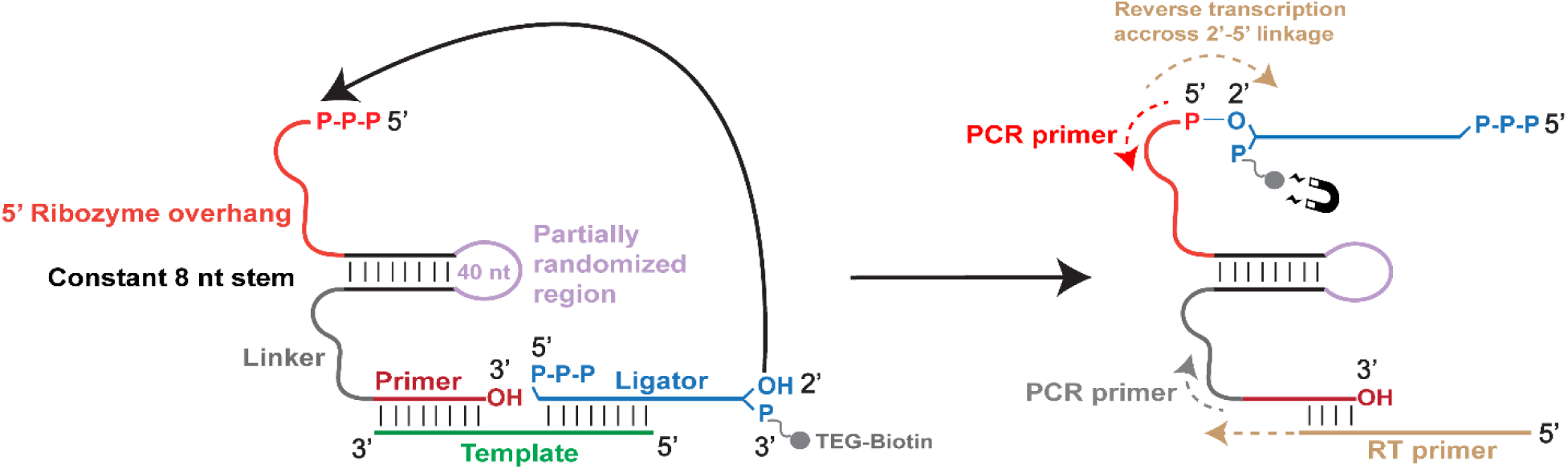
The unexpected ligation pathway exhibited by the isolated ribozymes. The ribozymes reported in this work catalyze ligation between the substrate 2′-hydroxyl and the ribozyme 5′-triphosphate groups. The 4 nt overlap between the RT primer (gold) and 3′ end of the ribozyme (‘primer’, red) allowed ligated products that were separated by streptavidin capture to be reverse transcribed. PCR primers targeting the 5′ and 3′ ends of the ribozyme amplified the ribozyme sequence to produce dsDNA as the template *for* in vitro transcription, which was used to generate the selection library for the subsequent round.

**Extended Data Fig. 9.**
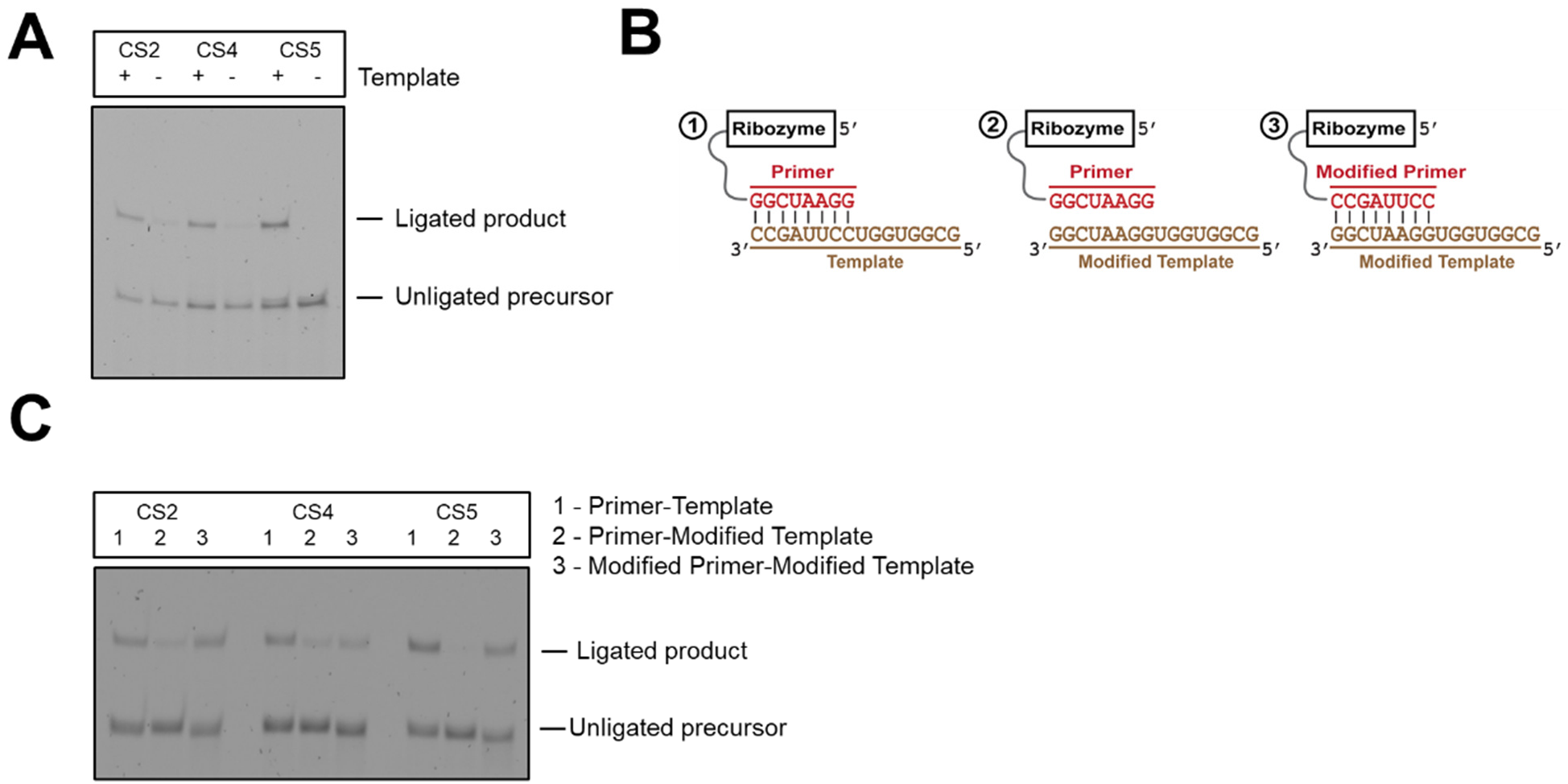
The role of template in RNA ligation by CS2, CS4, and CS5. A. CS2, CS4, and CS5 require an external template for ligation. B. Schematic for the compensatory mutational rescue of ligase activity in CS2, CS4, and CS5. C. Base-pairing interaction between the template and the 3′-primer sequence of CS2, CS4, and CS5 is important for ligation. Disrupting this interaction by mutations in the template abrogate ligation; however, compensatory mutations in the 3′-primer sequence of the ribozymes rescue ligation. Ligation reactions contained 1 µM ribozyme CS1, 1.2 μM RNA template, and 2 µM Substrate-3′P in 100 mM Tris-HCl (pH 8.0), 300 mM NaCl, and 100 mM MgCl_2_.

**Extended Data Fig. 10.**
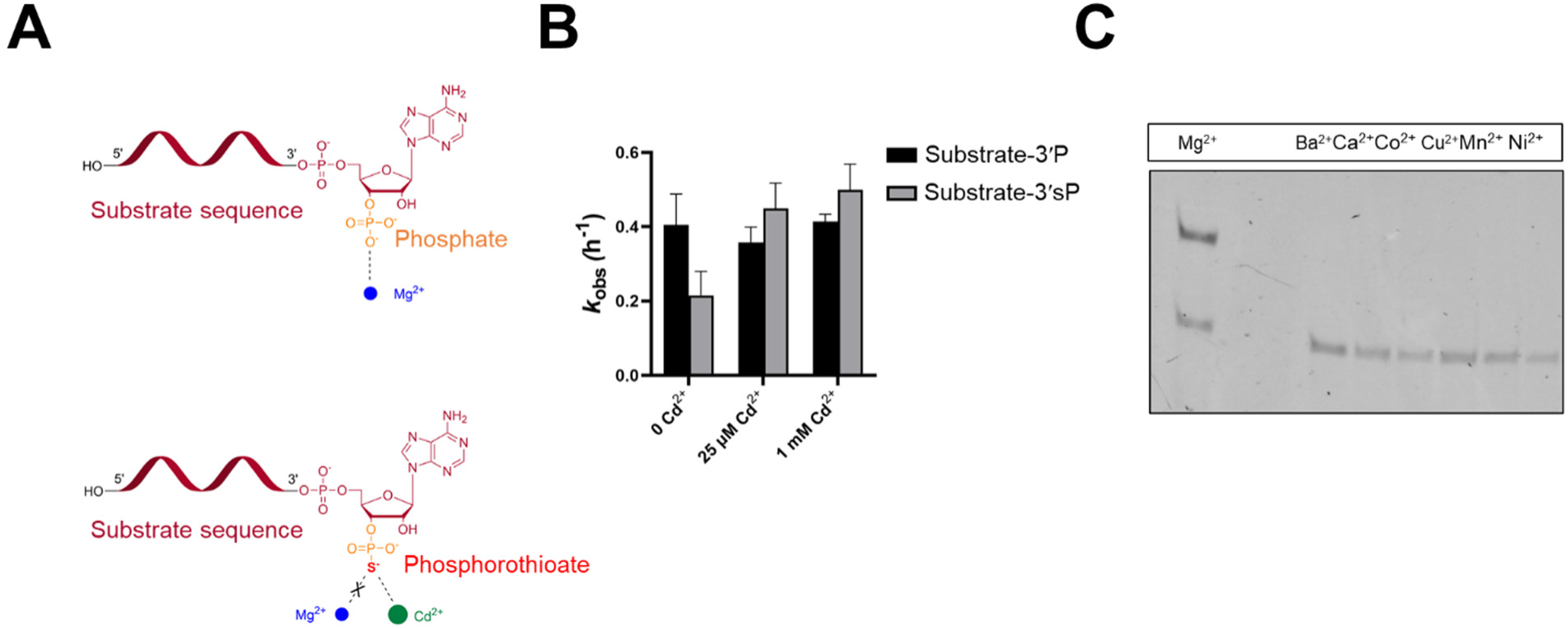
Mg^2+^ plays a potential catalytic role in ligation. A. Schematic illustrating a phosphorothioate metal rescue experiment. If the interaction between Mg^2+^ and a phosphate O is important for catalysis, replacing the O atom with a S atom (Substrate-3′sP), which weakens this interaction, is expected to cause a catalytic defect. Supplementing the reaction with thiophilic Cd^2+^ restores the putative interaction and is expected to rescue activity. B. A substrate with a 3′-thiophosphate causes a 2-fold reduction in ligation rate, which was rescued upon Cd^2+^ supplementation with an overall metal rescue of ∼2.5. C. Only Mg^2+^, among the divalent cations tested, supported ligation, suggesting a catalytic role. Ligation reactions contained 1 µM ribozyme and 2 µM substrate in 100 mM Tris-HCl (pH 8.0), 300 mM NaCl, 50 mM MgCl_2_ (B) or 100 mM MgCl_2_ (C), and the indicated amounts of Cd^2+^ (B) or 100 mM of the indicated divalent cations instead of Mg^2+^ (C).

**Extended Data Fig. 11.**
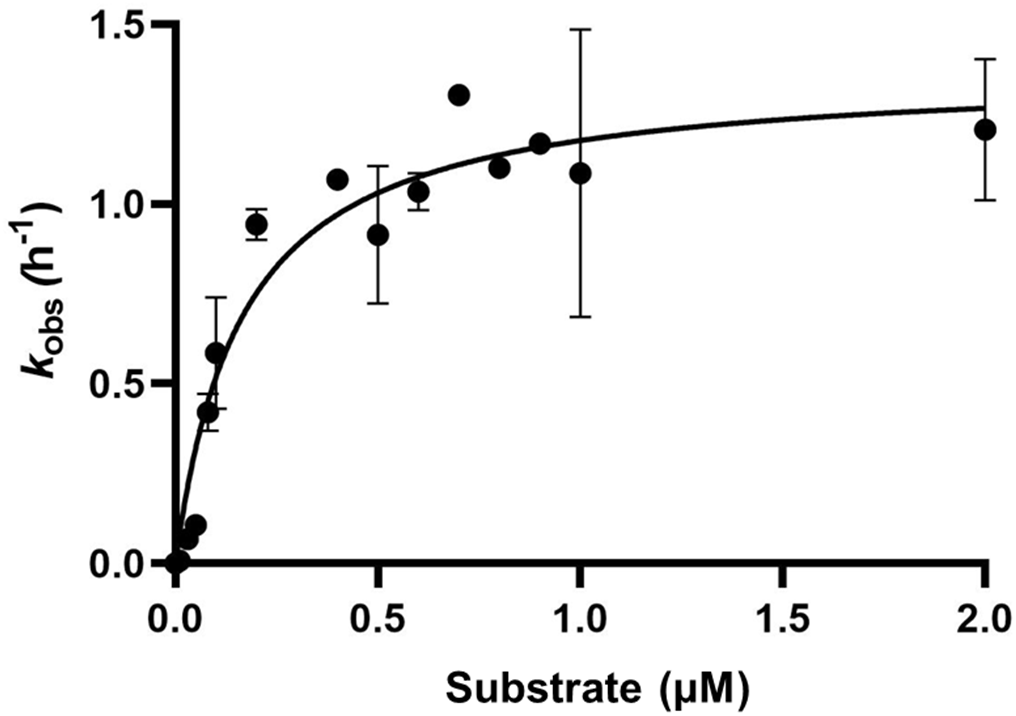
The effect of substrate concentration on reaction rates for CS1-catalyzed ligation. Michaelis-Menten plots for CS1-catalyzed ligation with Substrate-3′P reveal a *K*_M_ value of 0.1649 ± 0.042 µM and a *k*_cat_ / *K*_M_ value (catalytic efficiency) of 27.65 µM^-1^ h^-1^. Ligation reactions contained 1 µM ribozyme and the indicated amounts of Substrate-3′P in 100 mM Tris-HCl (pH 8.0), 300 mM NaCl, and 100 mM MgCl_2_.

**Extended Data Fig. 12.**
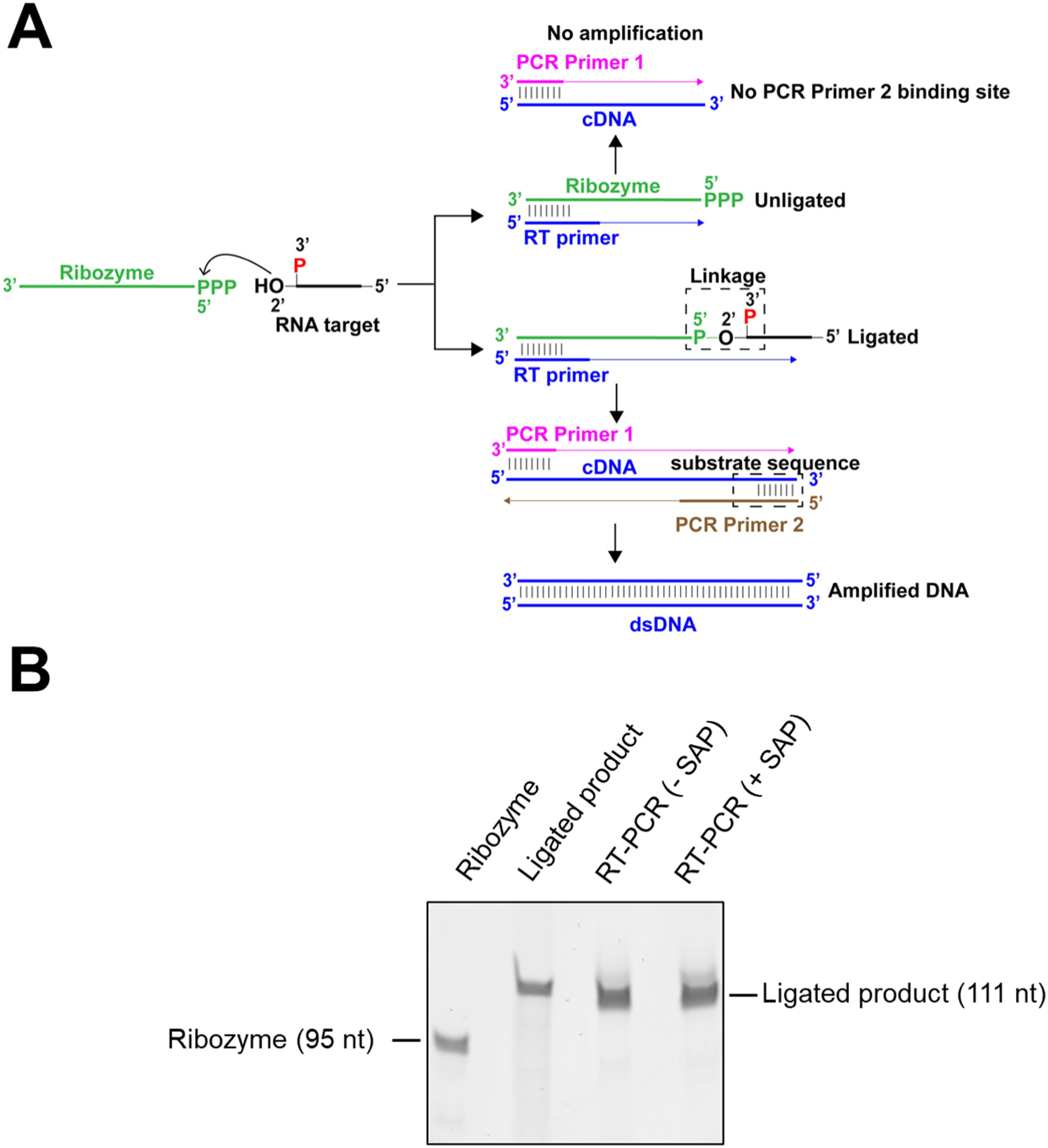
Ribozyme-assisted capture and amplification of 3′-phosphorylated RNA. A. Schematic illustrating the reverse transcription and PCR amplification of the ligated product generated by a reaction between CS1 and Substrate-3′P. B. Detection of a full-length dsDNA product after RT-PCR amplification of the ligated product, in the absence of a phosphate treatment step (-SAP) indicates successful reverse transcription across the unique 2′-5′, 3′P linkage between the ribozyme and the substrate. The SAP reaction was performed according to vendor specifications.

**Extended Data Fig. 13.**
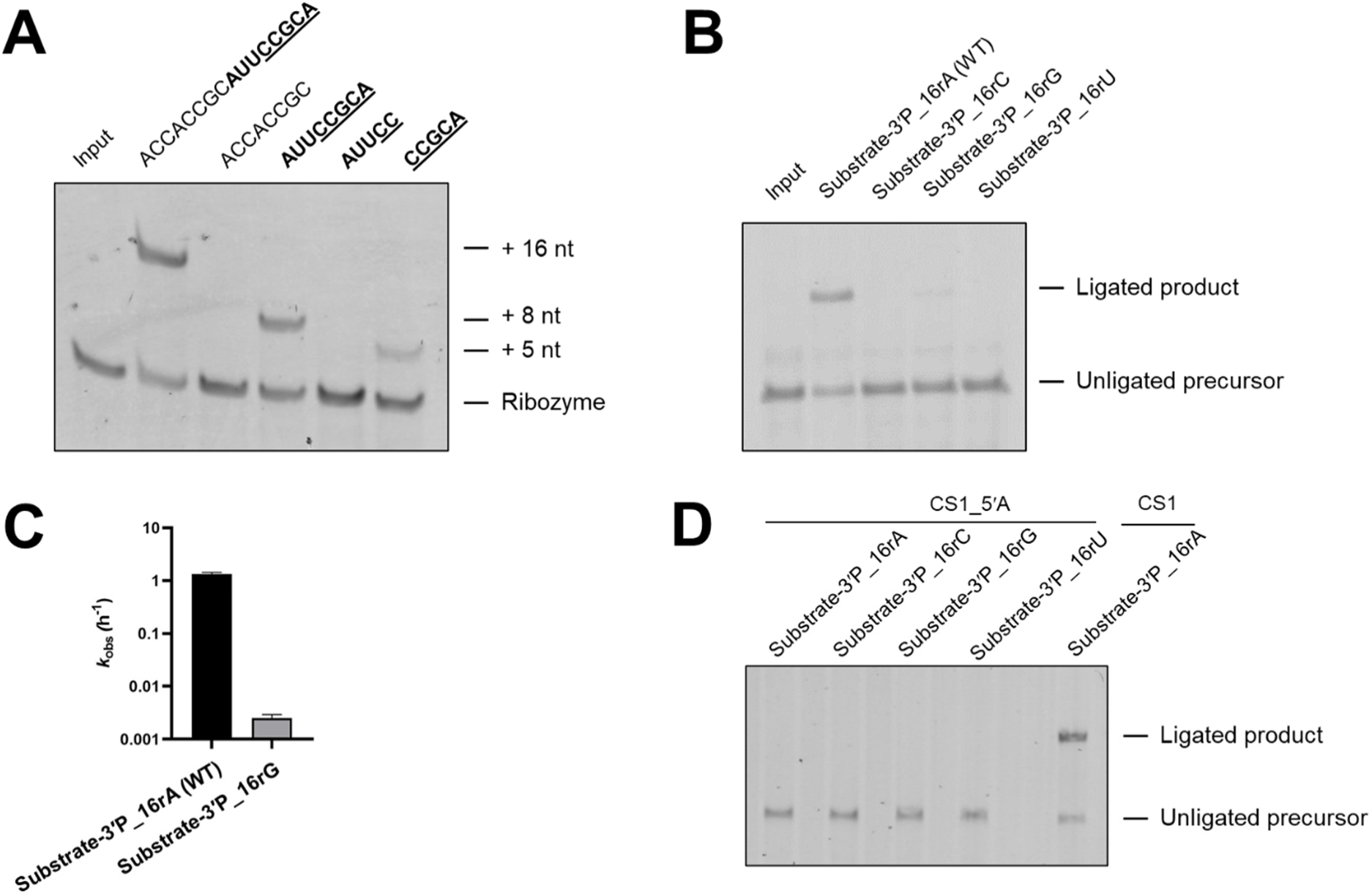
Substrate scope of ribozyme-catalyzed ligation. A. 3′-phosphorylated substrates of lengths ranging from 5-16 nt were active for ligation by CS1; however, only substrates containing the sequence CCGCA were ligated. B. The terminal adenine (the residue that contains the nucleophilic 2′-OH group) of the substrate is important for ligation. Substrates with terminal uridine or cytidine were inactive. Ligation with a substrate containing a terminal guanine was inefficient. C. Ligation was ∼500-fold slower with a substrate containing a terminal guanine relative to that with a substrate containing a terminal adenine. D. The 5′ terminal guanine (the residue that contains the electrophilic 5′-PPP group) of CS1 is indispensable for ligation. Ligation reactions contained 1 µM ribozyme and 2 µM substrate in 100 mM Tris-HCl (pH 8.0), 300 mM NaCl, and 100 mM MgCl_2_,

**Extended Data Fig. S14.**
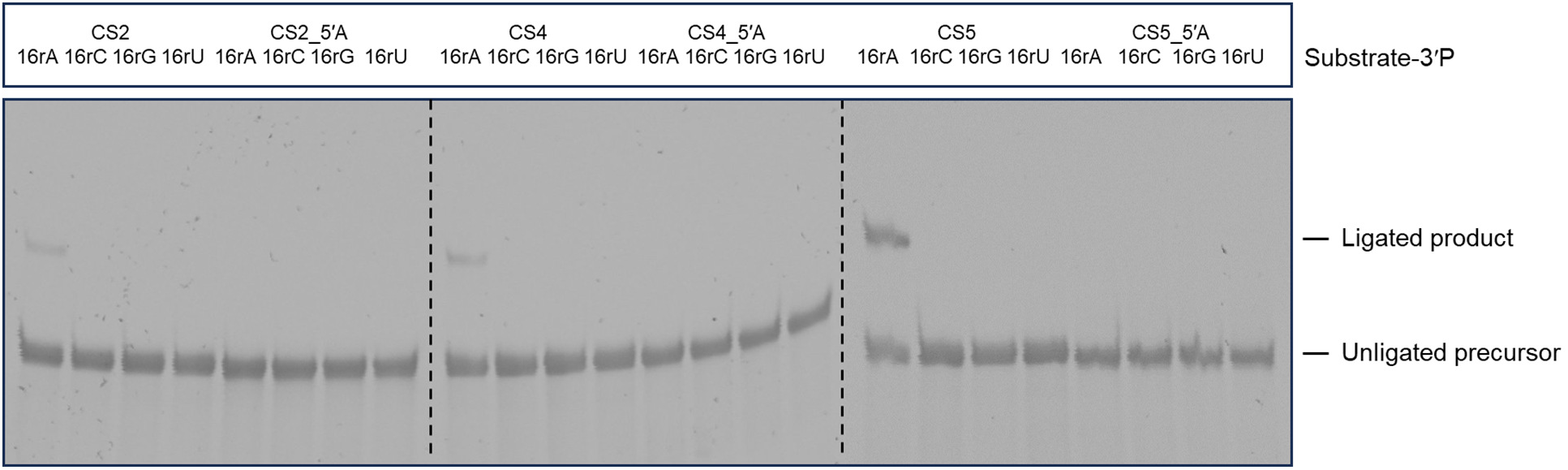
Sequence requirements for the ribozyme and substrate at the ligation junction for CS2, CS4, and CS5. The 5′ terminal guanine (the residue that contains the 5′-triphosphate group) of CS2, CS4, and CS5 and the terminal adenine (the residue that contains the nucleophilic 2′-hydroxyl group) of the substrate are essential for ligation. Ligation reactions contained 1 µM ribozyme, 1.2 µM template, and 2 µM substrate in 100 mM Tris-HCl (pH 8.0), 300 mM NaCl, and 100 mM MgCl_2_.

**Extended Data Fig. 15.**
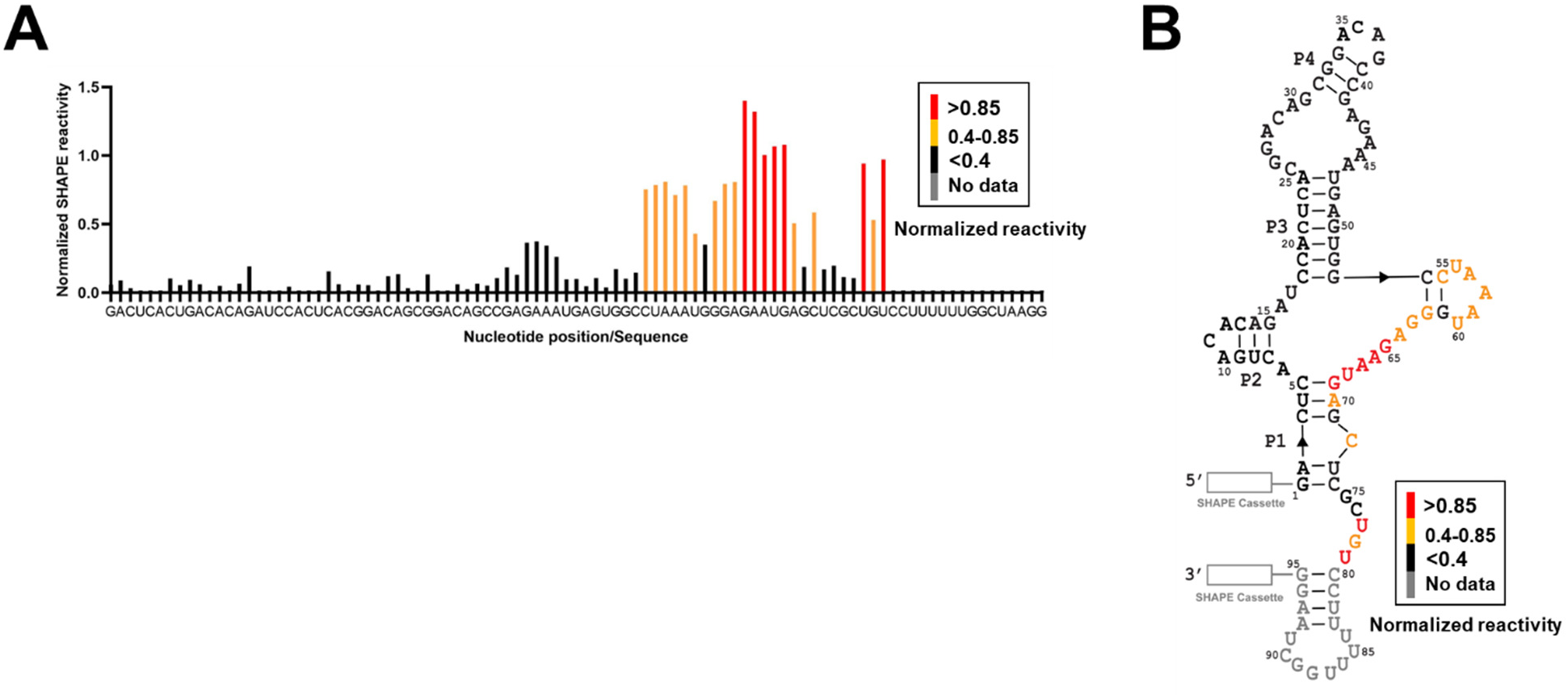
SHAPE analysis of CS1. Products of primer extension reactions with FAM-labeled primers after modification with 1M7 were separated on a 10% denaturing PAGE. A. Normalized SHAPE reactivities were plotted for each nucleotide position. High reactivity suggests flexibility within the RNA structure, while low reactivity indicates approximately base-paired stems or interactions with distal nucleotides in its tertiary fold. B. Secondary structure of CS1 determined by the RNAStructure program^36, 44^ using reactivity constraints obtained from SHAPE experiments. 5′ and 3′ SHAPE cassettes are denoted by white rectangles. Nucleotides for which no data was obtained are shown in gray. Nucleotides are colored in red, orange, and black according to their normalized SHAPE reactivities, as shown in the reactivity legend.

**Extended Data Fig. 16.**
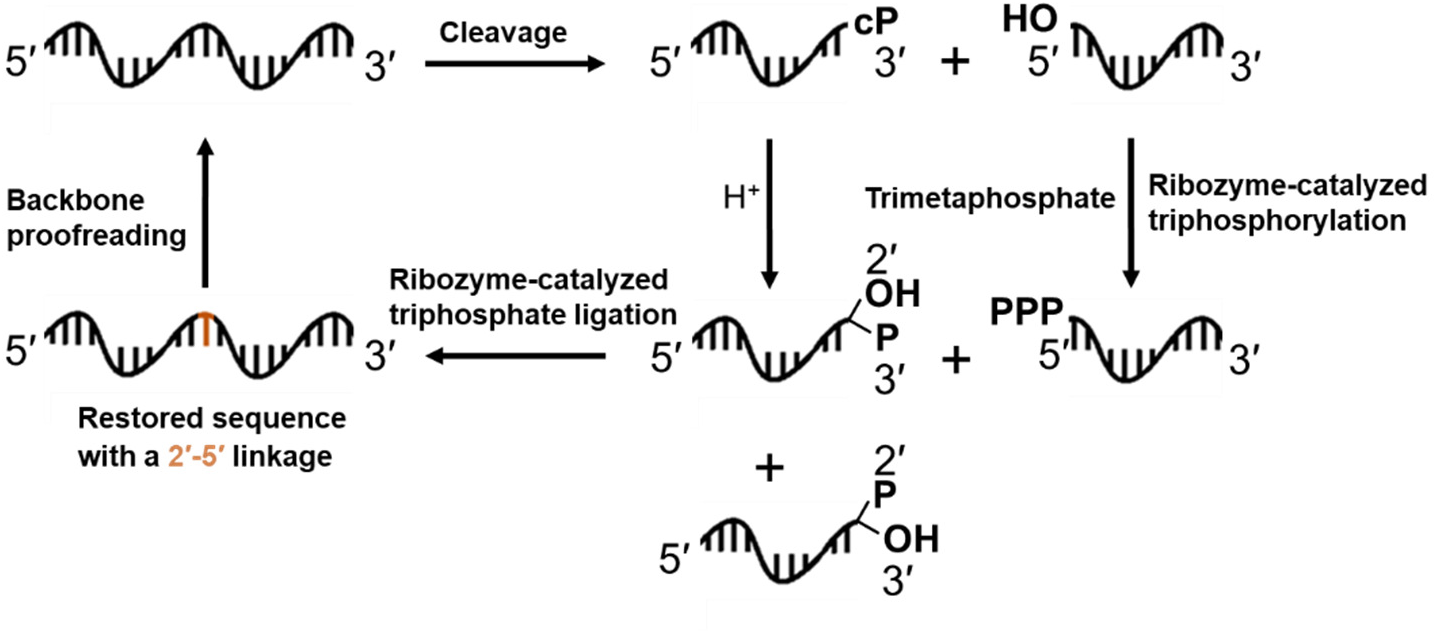
A hypothetical prebiotic pathway for ribozyme-assisted recovery of RNA cleavage products. The 5′ RNA cleavage product contains a 2′, 3′-cyclic phosphate is hydrolyzed to a 2′- or 3′-phosphate. The 5′ hydroxyl group of the 3′ cleavage product is activated with a triphosphate group by a ribozyme-catalyzed reaction that uses trimetaphosphate as the phosphorylating agent.^36^ A trans-acting ribozyme with the ligase activity reported in this work joins the resulting RNA oligonucleotides to regenerate the original RNA sequence pre-cleavage. This ligated product contains a 2′-5′ linkage in its backbone as opposed to an all 3′-5′ phosphodiester backbone in the original RNA. 2′-5′ linkages can be recycled nonenzymatically to 3′-5′ linkages through energy dissipative cycles.^41^

