## Supplementary Table 1 for "A ribozyme ligase that requires a 3′ terminal phosphate on its RNA substrate"

**Supplementary Table 1. Oligonucleotide sequences used in this work.** In the ribozyme sequences, variable nucleotides are highlighted in blue, the T7 promoter sequence is shown in purple, the hexauridine linker is shown in italics, and the 5′ terminal nucleotide possessing the triphosphate that participates in ligation is shown in boldface. 5′ and 3′ SHAPE cassettes are highlighted in green. Cleavage site nucleotides in the hairpin ribozyme substrate, FAM-HP_sub, are underlined. Oligonucleotides were either purchased from Integrated DNA Technologies (IDT) or Chemgenes or generated enzymatically by *in vitro* transcription (IVT) of dsDNA templates. AIP-Substrate was generated by incubating the corresponding 5′ monophosphorylated RNA with EDC and 2-aminoimidazole (2AI) (See 'RNA preparation and substrate activation' in Materials and Methods).

| # | Oligo Name | Sequence (5′→3′) | Type | Source |
|---|---|---|---|---|
| 1.1 | Parent AI ligase | GACUCACUGACACAGAUCCACUCACGGACAGCGGAAUGCUGCCAACCGUGCGGGCUAAUUGGCAGACUGAGCUCGCUGUCC*UUUUUU*GGCUAAGG | RNA | IVT |
| 1.2 | r0 DNA<br>(Mutagenesis at 21% at each nucleotide position: 79% WT nucleotide and 7% of the other three)<br>Note: The sequence is written according to IDT specifications | TAATACGACTCACTATA**G**ACTCACTGACACAGATCCACTCACGGACAGCG(N1:07077907)(N2:79070707)(N2)(N3:07070779)(N1)(N4:07790707)(N3)(N1)(N4)(N4)(N2)(N2)(N4)(N4)(N1)(N3)(N1)(N4)(N1)(N1)(N1)(N4)(N3)(N2)(N2)(N3)(N3)(N1)(N1)(N4)(N2)(N1)(N2)(N4)(N3)(N1)(N2)(N1)(N4)(N3)CGCTGTCC*TTTTTT*GGCTAAGG | DNA | IDT |
| 2.1 | Template | GCGGUGGUCCUUAGCC | RNA | IDT |
| 2.2 | Modified Template | GCGGUGGUGGAAUCGC | RNA | IDT |
| 3.1 | PPP-Substrate-Biot | (5′-triphosphate)-ACCACCGCAUUCCGCA -(3′-BioTEG) | RNA | Chemgenes |
| 3.2 | PPP-Substrate-diol | (5′-triphosphate)-ACCACCGCAUUCCGCA | RNA | Chemgenes |
| 3.3 | AIP- Substrate -Biot | (5′-phosphoro-2-aminoimidazole)-ACCACCGCAUUCCGCA-(3′-BioTEG) | RNA | Activation of P-Substrate-Biot |
| 3.4 | AIP-Substrate-diol | (5′-phosphoro-2-aminoimidazole)-ACCACCGCAUUCCGCA | | Activation of P-Substrate-diol |
| 3.5 | P- Substrate-Biot | (5′-monophosphate)-ACCACCGCAUUCCGCA-(3′-BioTEG) | RNA | IDT |
| 3.6 | Biot-P-Substrate-diol | (5BioTEG) (5′-monophosphate)-ACCACCGCAUUCCGCA | | |
| 3.7 | ddT-P- Substrate-Biot | (5′inverted ddT) (5′-monophosphate)-ACCACCGCAUUCCGCA-(3′-BioTEG) | RNA | IDT |
| 3.8 | Substrate-Biot (also, Substrate-TEG-Biot or HO-Substrate-Biot) | (5′-hydroxyl)-ACCACCGCAUUCCGCA-(3′-BioTEG) | RNA | IDT |
| 3.9 | Substrate-noTEG-Biot | (5′-hydroxyl)-ACCACCGCAUUCCGCA-(3′-Bio) | RNA | IDT |
| 3.10 | Substrate-TEG-DesthioBiot | (5′-hydroxyl)-ACCACCGCAUUCCGCA-(3′-deSBioTEG) | RNA | IDT |
| 3.11 | Substrate-3′P | (5′-hydroxyl)-ACCACCGCAUUCCGCA-(3′-monophosphate) | RNA | IDT |

| | | | | |
|---|---|---|---|---|
| 3.12 | FAM-Target RNA-3′P | (5′-FAM)- ACCACCGCAUUCCGCA-(3′-monophosphate) | RNA | IDT |
| 3.13 | Substrate-3′sP | (5′-hydroxyl)-ACCACCGCAUUCCGCA-(3′-monothiophosphate) | RNA | IDT |
| 3.14 | P-Substrate16dA-Biot | (5′-monophosphate)-ACCACCGCAUUCCGCdA-(3′-BioTEG) | RNA | IDT |
| 3.15 | Substrate-3′P_8mer1 | (5′-hydroxyl)-ACCACCGC-(3′-monophosphate) | RNA | IDT |
| 3.16 | Substrate-3′P_8mer2 | (5′-hydroxyl)-AUUCCGCA-(3′-monophosphate) | RNA | IDT |
| 3.17 | Substrate-3′P_5mer1 | (5′-hydroxyl)-AUUCC-(3′-monophosphate) | RNA | IDT |
| 3.18 | Substrate-3′P_5mer2 | (5′-hydroxyl)-CCGCA-(3′-monophosphate) | RNA | IDT |
| 3.19 | Substrate-3′P_16rA | (5′-hydroxyl)-ACCACCGCAUUCCGCA-(3′-monophosphate) | RNA | IDT |
| 3.20 | Substrate-3′P_16rC | (5′-hydroxyl)-ACCACCGCAUUCCGCC-(3′-monophosphate) | RNA | IDT |
| 3.21 | Substrate-3′P_16rG | (5′-hydroxyl)-ACCACCGCAUUCCGCG-(3′-monophosphate) | RNA | IDT |
| 3.22 | Substrate-3′P_16rU | (5′-hydroxyl)-ACCACCGCAUUCCGCU-(3′-monophosphate) | RNA | IDT |
| 4.1 | RT primer | GTGCGGAATGCGGTGGTCCTT | DNA | IDT |
| 4.2 | SHAPE_RT_primer | (5′-FAM)-GAACCGGACCGAAGCCCG | DNA | IDT |
| 5.1 | PCR_Fwd_primer | TAATACGACTCACTATAGACTCACTGACAC | DNA | IDT |
| 5.2 | PCR_LigFwd_primer | ACCACCGCATTCCG | DNA | IDT |
| 5.3 | PCR_Rvs_primer | mCmCTTAGCCAAAAAAGGACAGCG | DNA | IDT |
| 6.1 | CS1 | **G**ACUCACUGACACAGAUCCACUCACGGACAGCGGACAGCCGAGAAAUGAGUGGCCUAAAUGGGAGAAUGAGCUCGCUGUCC*UUUUUU*GGCUAAGG | RNA | IVT |
| 6.2 | CS1_5’A | **A**ACUCACUGACACAGAUCCACUCACGGACAGCGGACAGCCGAGAAAUGAGUGGCCUAAAUGGGAGAAUGAGCUCGCUGUCC*UUUUUU*GGCUAAGG | RNA | IVT |
| 6.3 | CS1_5′ truncated | GGACAGCGGACAGCCGAGAAAUGAGUGGCCUAAAUGGGAGAAUGAGCUCGCUGUCC*UUUUUU*GGCUAAGG | RNA | IVT |
| 6.4 | CS1_3′ truncated | **G**ACUCACUGACACAGAUCCACUCACGGACAGCGGACAGCCGAGAAAUGAGUGGCCUAAAUGGGAGAAUGAGCUCGCUGUCC | RNA | IVT |
| 6.5 | CS1_5′+3′ truncated | GGACAGCGGACAGCCGAGAAAUGAGUGGCCUAAAUGGGAGAAUGAGCUCGCUGUCC | RNA | IVT |
| 6.6 | CS1_SHAPE | GGCCUUCGGGCCAA**G**ACUCACUGACACAGAUCCACUCACGGACAGCGGACAGCCGAGAAAUGAGUGGCCUAAAUGGGAGAAUGAGCUCGCUGUCCUUUUUUGGCUAAGGUCGAUCCGGUUCGCCGGAUCCAAAUCGGGCUUCGGUCCGGUUC | RNA | IVT |
| 6.7 | CS2 | **G**ACUCACUGACACAGAUCCACUCACGGACAGCGGACUGCGCGUAUGAGUGGCGGCUAAAGAGGAGAAUGAGCGCGCUGUCC*UUUUUU*GGCUAAGG | RNA | IVT |
| 6.8 | CS2_5’A | **A**ACUCACUGACACAGAUCCACUCACGGACAGCGGACUGCGCGUAUGAGUGGCGGCUAAAGAGGAGAAUGAGCGCGCUGUCC*UUUUUU*GGCUAAGG | RNA | IVT |
| 6.9 | CS2_modified primer | **G**ACUCACUGACACAGAUCCACUCACGGACAGCGGACUGCGCGUAUGAGUGGCGGCUAAAGAGGAGAAUGAGCGCGCUGUCC*UUUUUU*CCGAUUCC | RNA | IVT |

| | | | | |
|---|---|---|---|---|
| 6.10 | CS3 | GACUCACUGACACAGAUCCACUCACGGACAGC GACGGGUGGGUAAUCUAGUGUCCGCGGAAUAG AACGAAACACGCUGUCC*UUUUUU*GGCUAAGG | RNA | IVT |
| 6.11 | CS4 | **G**ACUCACUGACACAGAUCCACUCACGGACAGC GGGAUGGUGCGAACUGAGUGGGCUAAUUAGGA GAAUGAGCGCGCUGUCC*UUUUUU*GGCUAAGG | RNA | IVT |
| 6.12 | CS4_5'A | **A**ACUCACUGACACAGAUCCACUCACGGACAGC GGGAUGGUGCGAACUGAGUGGGCUAAUUAGGA GAAUGAGCGCGCUGUCC*UUUUUU*GGCUAAGG | RNA | IVT |
| 6.13 | CS4_modified primer | **G**ACUCACUGACACAGAUCCACUCACGGACAGC GGGAUGGUGCGAACUGAGUGGGCUAAUUAGGA GAAUGAGCGCGCUGUCC*UUUUUU*CCGAUUCC | | |
| 6.14 | CS5 | **G**ACUCACUGACACAGAUCCACUCACGGACAGC GGGAGGGUGACAUCGUUGAGAGAGAAUGGGGA UAUUGAACUCGCUGUCC*UUUUUU*GGCUAAGG | RNA | IVT |
| 6.15 | CS5_5'A | **A**ACUCACUGACACAGAUCCACUCACGGACAGC GGGAGGGUGACAUCGUUGAGAGAGAAUGGGGA UAUUGAACUCGCUGUCC*UUUUUU*GGCUAAGG | RNA | IVT |
| 6.16 | CS5_modified primer | **G**ACUCACUGACACAGAUCCACUCACGGACAGC GGGAGGGUGACAUCGUUGAGAGAGAAUGGGGA UAUUGAACUCGCUGUCC*UUUUUU*CCGAUUCC | RNA | IVT |
| 7.1 | PPP-CS1_pc1 | (5'-triphosphate)- GACUCACUGACACAGAUCCACUCAC | RNA | IVT |
| 7.2 | P-CS1_pc1 | (5'-monophosphate)- GACUCACUGACACAGAUCCACUCAC | RNA | IDT |
| 7.3 | HO-CS1_pc1 | (5'-hydroxyl)- GACUCACUGACACAGAUCCACUCAC | RNA | IDT |
| 7.4 | pCS1_pc2 | (5'-monophosphate)- GGACAGCGGACAGCCGAGAAAUGAGUGGCCUA AAUGGGAG | RNA | IDT |
| 7.5 | pCS1_pc3 | (5'-monophosphate)- AAUGAGCUCGCUGUCCUUUUUUGGCUAAGG | RNA | IDT |
| 7.6 | CS1_splint1 | CTCGGCTGTCCGCTGTCCGTGAGTGGATCTGT GTCAG | DNA | IDT |
| 7.7 | CS1_splint2 | CCAAAAAAGGACAGCGAGCTCATTCTCCCATT TAGGCCAC | DNA | IDT |
| 8.1 | FAM-HP_sub | (5'-FAM)- ACCACCGCAUUCCGCAGUCCUCUCC | RNA | IDT |
| 8.2 | HP_ribozyme | GGAGAGAGAAGCGGACCAGAGAAACACACGUU GUGGUAUAUUACCUGGUA | RNA | IVT |
